# Genotype Representation Graphs: Enabling Efficient Analysis of Biobank-Scale Data

**DOI:** 10.1101/2024.04.23.590800

**Authors:** Drew DeHaas, Ziqing Pan, Xinzhu Wei

**Affiliations:** Department of Computational Biology, Cornell University, Ithaca, NY

**Author notes:** Contributed equally (Drew DeHaas, Ziqing Pan).

## Abstract

Computational analysis of a large number of genomes requires a data structure that can represent the dataset compactly while also enabling efficient operations on variants and samples. Current practice is to store large-scale genetic polymorphism data using tabular data structures and file formats, where rows and columns represent samples and genetic variants. However, encoding genetic data in such formats has become unsustainable. For example, the UK Biobank polymorphism data of 200,000 phased whole genomes has exceeded 350 terabytes (TB) in Variant Call Format (VCF), cumbersome and inefficient to work with. To mitigate the computational burden, we introduce the Genotype Representation Graph (GRG), an extremely compact data structure to losslessly present phased whole-genome polymorphisms. A GRG is a fully connected hierarchical graph that exploits variant-sharing across samples, leveraging ideas inspired by Ancestral Recombination Graphs. Capturing variant-sharing in a multitree structure compresses biobank-scale human data to the point where it can fit in a typical server’s RAM (5-26 gigabytes (GB) per chromosome), and enables graph-traversal algorithms to trivially reuse computed values, both of which can significantly reduce computation time. We have developed a command-line tool and a library usable via both C++ and Python for constructing and processing GRG files which scales to a million whole genomes. It takes 160GB disk space to encode the information in 200,000 UK Biobank phased whole genomes as a GRG, more than 13 times smaller than the size of compressed VCF. We show that summaries of genetic variants such as allele frequency and association effect can be computed on GRG via graph traversal that runs significantly faster than all tested alternatives, including *vcf.gz*, PLINK BED, tree sequence, XSI, and Savvy. Furthermore, GRG is particularly suitable for doing repeated calculations and interactive data analysis. We anticipate that GRG-based algorithms will improve the scalability of various types of computation and generally lower the cost of analyzing large genomic datasets.

## Introduction

The rapid progress in sequencing technologies and the growing interest in genetic association with diseases have led to the collection of large datasets comprising hundreds of thousands of sequenced human whole genomes (Kaiser 2021). At the end of 2023, the UK Biobank released ∼500,000 whole genomes on their cloud computing platform, 200,000 of which have already been phased (Browning and Browning 2023; Hofmeister et al., 2023). To store a large number of phased whole genomes, the formats VCF (sometimes called pVCF, Danecek et al., 2011, **Fig. 1a**), and BGEN (Band and Marchini, 2018) are the most commonly used. These two formats vary in whether they store genotypes as hard calls (i.e., variant allele or not a variant) or probabilities of being a variant, and whether they use text or binary format. Despite the differences, genetic polymorphism data are generally organized either by individuals or by variants, to be easily converted into a 2-D matrix (**Fig. 1b**) or a 1-D vector when being loaded into computer memory for downstream computations. We hereafter refer to such formats as tabular formats.

**Figure 1:**
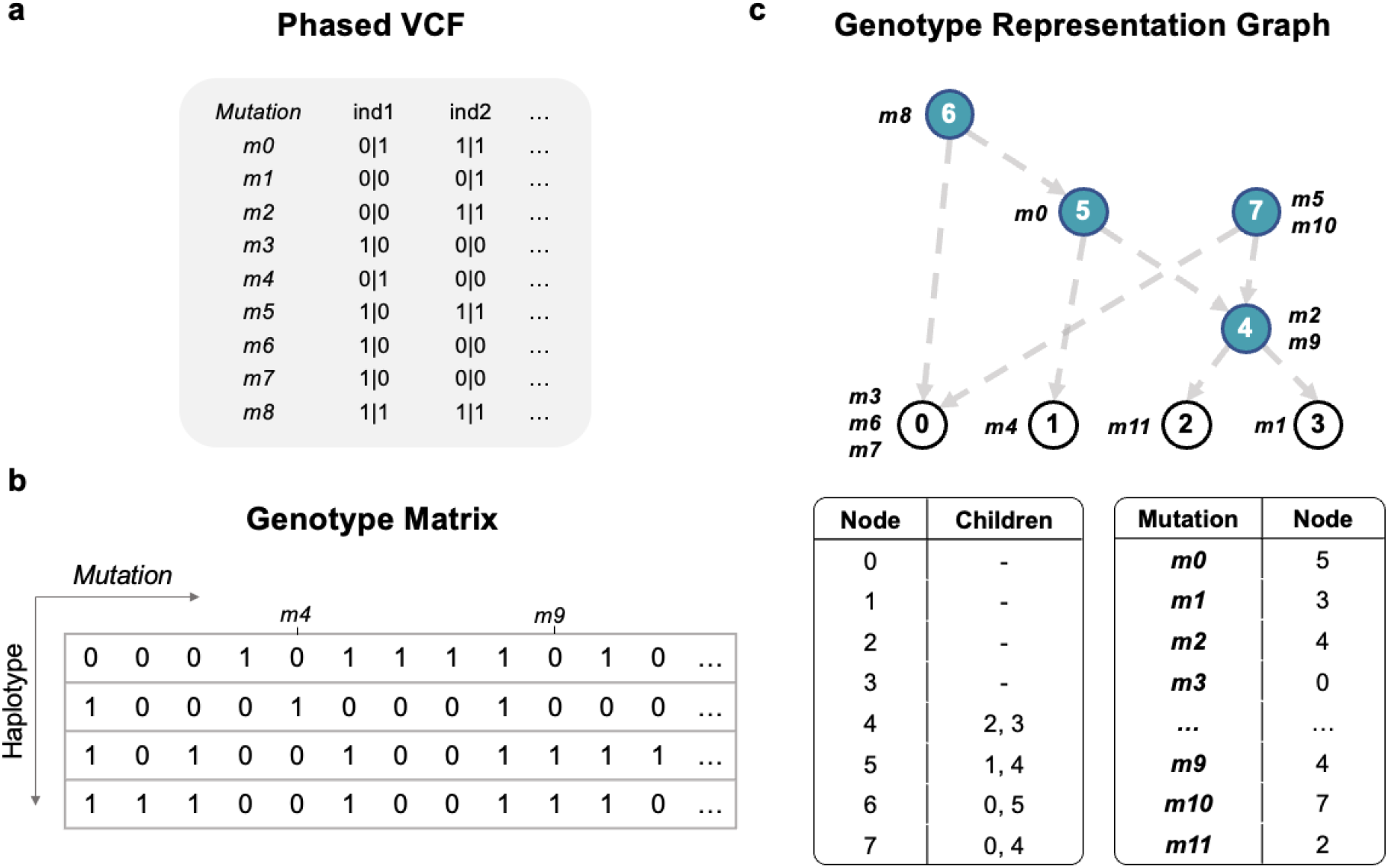
Data encodings for phased polymorphisms. Phased diploid genotypes in three different formats. **a**: A phased VCF file. Rows are genetic polymorphisms, columns are individual genotypes. The reference allele is labeled as 0 and alternative alleles are non-zero. A vertical bar separates an individual’s genotype into two haplotypes. **b**: Encoding the data as a matrix. This is essentially a rotation of the VCF data, with columns representing polymorphisms and rows representing haplotypes. **c**: A GRG representation of the same data. Each node represents a unique subset of samples, and each mutation is mapped to one node. The node IDs are labeled inside nodes, and the mutation IDs are marked next to the mutation nodes. Samples are reachable from a mutation node iff the mutation is present in the sample haplotype. This GRG can be described as the tables in the lower panel **c**.

The convenience and ease of use of tabular data structures have made them the predominant and default data structures for encoding genotypes. However, encoding biobank-scale whole-genome polymorphisms with uncompressed conventional tabular formats has become unsustainable. In addition to VCF and BGEN, some tools and libraries make use of other tabular formats like BCF (Li 2011) and BED (PLINK) (Purcell et al., 2007). Recent research on genetic file formats has led to the development of many other tabular formats such as GTC (Danek and Deorowicz 2018), GQT (Layer et al., 2016), bref3 (Browning et al., 2018), genozip (Lan et al., 2021), GTSshark (Deorowicz and Danek 2019), PGEN (PLINK2) (Rivas and Chang 2024), XSI (Wertenbroek et al., 2022), and Savvy (LeFaive et al., 2021). Among these, the formats that achieve the best compression (such as GTShark, XSI, and Savvy) are based on positional Burrows-Wheeler transform (PBWT) (Durbin 2014). PBWT imposes a sorted order of samples that captures shared haplotypes between them, and this can help compress the data in conjunction with encodings (e.g., word-aligned hybrid (Wu 2001)). Additionally, many of these formats take advantage of standard compression algorithms such as zstd (Collet and Kucherawy 2021) to further compress the data.

While exploiting the compressed sparse data representations alleviates the burden of large-scale data encodings, it introduces additional computational cost for downstream analysis due to the need to decompress the data. To address this issue, a desirable data structure for population genotypes should mimic the true underlying biological generative model of DNA sequences, representing the process in a compact but uncompressed form. Notably, the generative model of DNA sequences has been studied in population genetics for decades. In the absence of incomplete lineage sorting (Kapli et al., 2020), the genetic differences among species generated by spontaneous mutations can be seen as sampling random mutations based on the mutation rate on each branch of a true phylogenetic tree. Therefore, assuming DNA sequences can be accurately ascertained, a phylogenetic tree can be inferred from DNA sequences at the species level (Fitch and Margoliash, 1967; Kapli et al., 2020), and these sequences can be efficiently stored with a tree data structure (**Fig. S1**). In the absence of recombination, a tree structure fully describes the evolutionary relationship among a set of genomes. In the presence of recombination, different sections of the genomes trace back to different common ancestors. As a result, the evolutionary relationship among a set of genomes can no longer be described by a single tree structure, but a graph. A graph of sequences of closely related species is called a phylogenetic network (Huson and Bryant, 2006), and a graph of sequences within species is called an Ancestral Recombination Graph (ARG) (Brandt et al., 2022; Brandt et al., 2024; Lewanski et al., 2024; Wong et al., 2024). Considering that individual haplotypes in a population are related through true genealogical history (Hudson 1983a), the genetic variants among haplotypes that arise from spontaneous mutations can be seen as a random sampling process on the ARG (Tavaré 1984; Zhang et al., 2023). When an ARG is inferred from a set of *phased* whole genomes, the resulting graph can be used to store information about these genomes. However, mutations only occur very sparsely on an ARG, so using an ARG to encode genetic variants for downstream computations might not be efficient. Moreover, ARG inference for more than 10,000 whole genomes is currently computationally prohibitive (Rasmussen et al., 2014; Speidel et al., 2019; Kelleher et al., 2019; Brandt et al., 2022; Deng et al., 2024). As a result, ARG data structures have not been adopted for biobank-scale whole-genome statistical analyses.

In this study, we introduce the Genotype Representation Graph (GRG) (**Fig. 1c**), a graph representation of whole genomes, and implement a scalable method that can construct a GRG for a million whole genomes at a very low cost. GRG is designed to faithfully encode the hard-call information in phased whole genomes, while compressing the data significantly. Moreover, a GRG can be traversed efficiently which enables algorithms in the style of dynamic programming (Eddy, 2004), facilitating scalable genome-wide statistical analyses. GRG occupies the design space between ARGs and PBWT-based compression formats like XSI and Savvy. Specifically, GRG leverages the shared hierarchies that represent shared haplotype structures among samples to compress the data. To our knowledge, it is the first hierarchical genotype compression format, and is competitive with the best formats for file size and scalability.

## Results

### Overview of GRG

We use the term *mutation* to refer to a (genomic position, derived allele) pair, such that each genomic position could have more than one mutation. In practice, we assume no recurrent mutations, and the derived alleles are often approximated with non-reference or minor alleles. In general, a single haplotype can be represented as the full DNA sequence or as a sparse list of mutations. Such a list contains only the variations from the reference sequence: a haplotype that exactly matches the reference at all polymorphic sites is represented by an empty list. Similarly, a GRG explicitly encodes phased haplotypes as lists of mutations: only deviations from a reference sequence are represented in the graph.

A GRG is a directed acyclic graph (DAG) (Greenland et al., 1999) with the following properties:

I. The nodes with no successors (leaf nodes) are *sample nodes*. There is one sample node for each haploid genome in the dataset, and a diploid individual will be represented by two sample nodes.
II. Any node (including sample nodes) can contain zero or more mutations. A node containing at least one mutation is called a *mutation node*. A node containing zero mutations is called an *empty node*.
III. A node that has at least one successor and one predecessor is called an *internal node*. An internal node represents the union of samples beneath it, and the union of mutations above it. We say such a node *covers* these samples and mutations.
IV. The roots (nodes with no predecessors) must be mutation nodes.
V. There exists at most one path between any two nodes (i.e., a multitree structure).
VI. There exists a path from the mutation node containing mutation *M*_i_ to the sample node *S*_j_ iff the genotype for sample *S*_j_ contains mutation *M*_i_.

A GRG is defined by its *down edges*: the edges that are directed from the roots to the samples. We also refer to this as the parent-child direction. It also symmetrically contains *up edges*, which are child-parent edges and form the reverse DAG.

Attached to each GRG node is a count of the number of diploid individuals whose two sample nodes coalesce at that node. This information is required for certain statistical calculations (such as genome-wide association study (GWAS)).

The worst-case GRG is equivalent to representing the dataset as a list of mutations: each mutation node has a direct edge to every sample that contains it. In contrast, a compact GRG utilizes internal nodes to cover sample subsets that are shared by many mutations, thus compressing the mutation-to-sample relationship (see **Fig. S2**).

### A Scalable Algorithm for Constructing GRGs

The goal of GRG construction is to create a graph representation of the relationship among samples such that 1) each mutation is correctly contained in a node and 2) the number of edges in the graph is minimized. To this end, GRG construction proceeds in four high-level steps (**Fig. 2a**). The first step can be conceptualized as separating the genotype matrix into *T* non-overlapping continuous segments. For each of these segments, a local GRG is constructed through two additional steps: (2) building a tree GRG (tGRG) that approximates the genetic relationship among all samples based on the genotypes in the current segment (the *BuildShape* step), and (3) mapping mutations from the current segment to the tGRG (the *MapMutation* step). Finally, the *T* tGRGs are merged into a single global GRG that represents the genome-wide polymorphisms. Experimentally, it is optimal to split the genome into segments of size 50-150Kbp (kilo base pairs) for most non-simulated human datasets (**Fig. S3**). The final GRG is then simplified and serialized to disk based on its down edges (see **Supplement**), but both down and up edges can be kept in RAM to facilitate a wide variety of computations.

**Figure 2:**
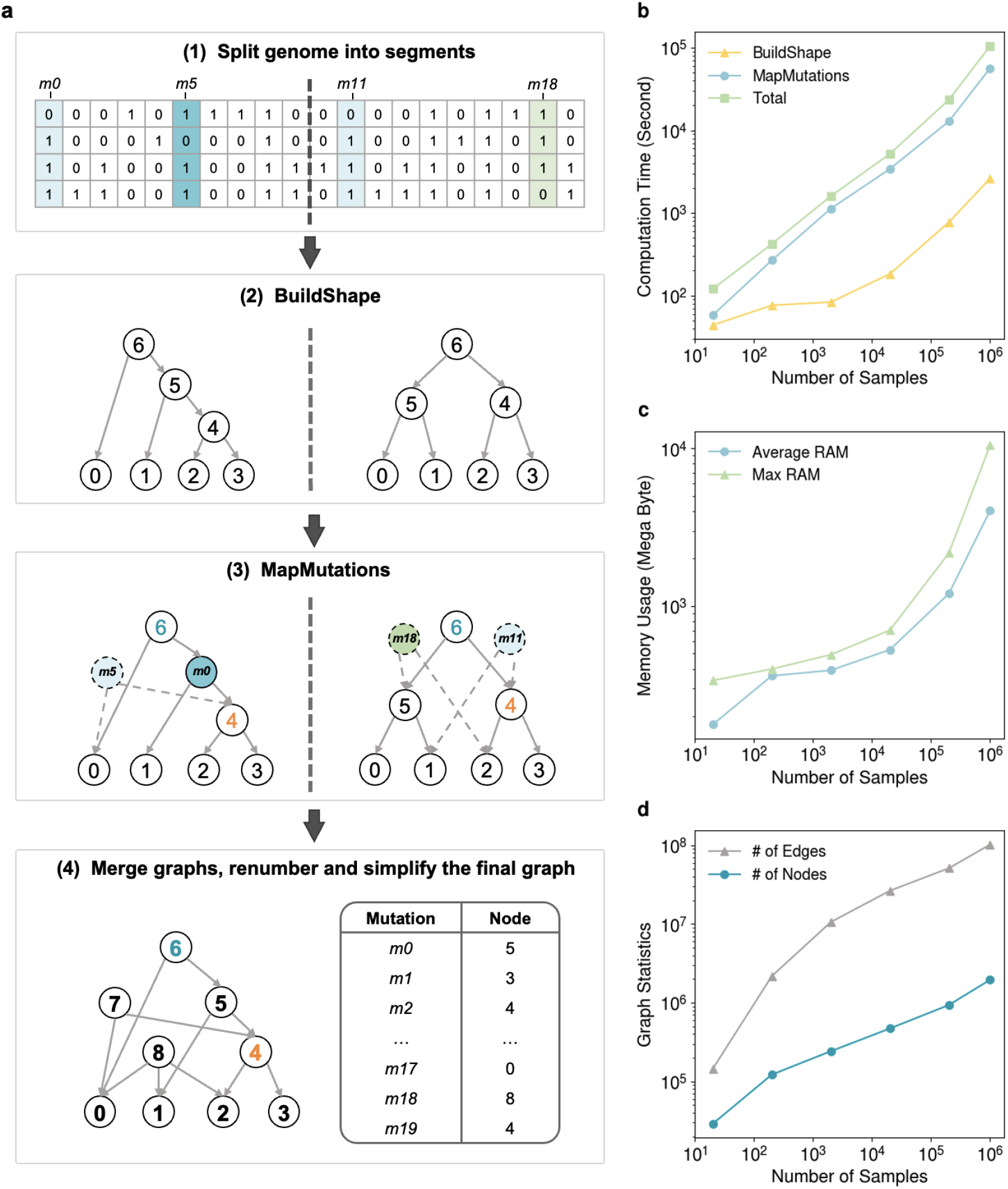
Overview and performance of the GRG construction algorithm. **a**: GRG construction includes four high-level steps: (1) segmenting the genome, (2) building an initial tGRG for each segment, (3) mapping mutations from each segment to the corresponding tGRG, and (4) merging all tGRGs into a single GRG. Dashed lines represent edges/nodes newly created in the current step. **b, c**, and **d** summarize the performance of GRG construction against various sample sizes using simulated data of a 100 mega base pair (Mbp) region. **b**: The total construction time (in green) and the time breaking down into BuildShape (in yellow) and MapMutations (in blue) against the number of samples. **c**: Memory usage in terms of average RAM (in blue) and maximum RAM (in green) against the number of samples. **d**: Graph size of the resulting GRGs breaking down into the number of edges (in grey) and number of nodes (in teal) against the number of samples.

The *BuildShape* and *MapMutations* steps are tightly linked. When a mutation’s sample list can be fully represented by an existing node in the graph, the mutation is directly mapped to the node (e.g., mutation *m0* in **Fig. 2a**). Otherwise, a new node is created and connected to existing nodes in the graph such that the union of samples of the new node can exactly represent the sample list of the mutation (e.g., mutation *m5* in **Fig. 2a**). In the worst case, a new node is added for the mutation, and edges are added from the new node to all samples that contain the mutation. *BuildShape* is designed to avoid this worst-case scenario by generating nodes that are highly reusable across many mutations.

In each segment separated in step (1), *BuildShape* creates a tGRG by repeatedly modifying and merging pairs of subtrees from a set until a single binary tree is formed. The initial set is simply the sample nodes, serving as the roots of the subtrees. An arbitrary root node from the subtree set is selected, and a nearest-neighbor search is performed to find another root node of the highest similarity measured by the Hamming distance (Waggener, 1995) of the mutation lists associated with the nodes. The Hamming distance is against the mutation lists (haplotypes) for the associated samples, when the sample nodes are root nodes. As new root nodes are added, Hamming distance is against the consensus haplotype (Miller et al., 2010) represented by the root, i.e., the intersection of their descendants’ mutation lists. After the nearest-neighbor search, the two subtrees are merged by constructing a new root node and attaching the previous root nodes to it. These steps are repeated until a single tree is formed (only one root node remains in the set), which is the tGRG. The mutations are then mapped, either directly to the tGRG or to a GRG composed of multiple tGRGs that have been merged. During mutation mapping, each node’s individual coalescence count is stored, since the list of samples beneath each node is already being traversed.

### GRG Construction Results

We first tested the performance characteristics of GRG construction using simulated data in terms of computation time (**Fig. 2b**), memory usage (**Fig. 2c**), and graph statistics (**Fig. 2d**). We used *msprime* (Kelleher et al., 2016; Baumdicker et al., 2022) to simulate 100Mbp sequences for up to 500,000 diploid individuals (1,000,000 haploid samples) with a human-like parameter setting (Ne = 10^4^, recombination rate = 10^−8^ / (bp × generation), mutation rate = 10^−8^ / (bp × generation)). GRG construction is memory efficient, with a maximum RAM usage of 10GB for a million samples (**Fig. 2c**). The file size of the resulting GRGs are shown in **Fig. 3a**. In addition to the standard formats .*vcf*.*gz* and BED, we compare GRG against XSI and Savvy. Of the various compression formats, XSI and Savvy provide some of the best compression rates (Wertenbroek et al., 2022), and also provide APIs for performing computations on the data without having to decompress the entire file first.

**Figure 3:**
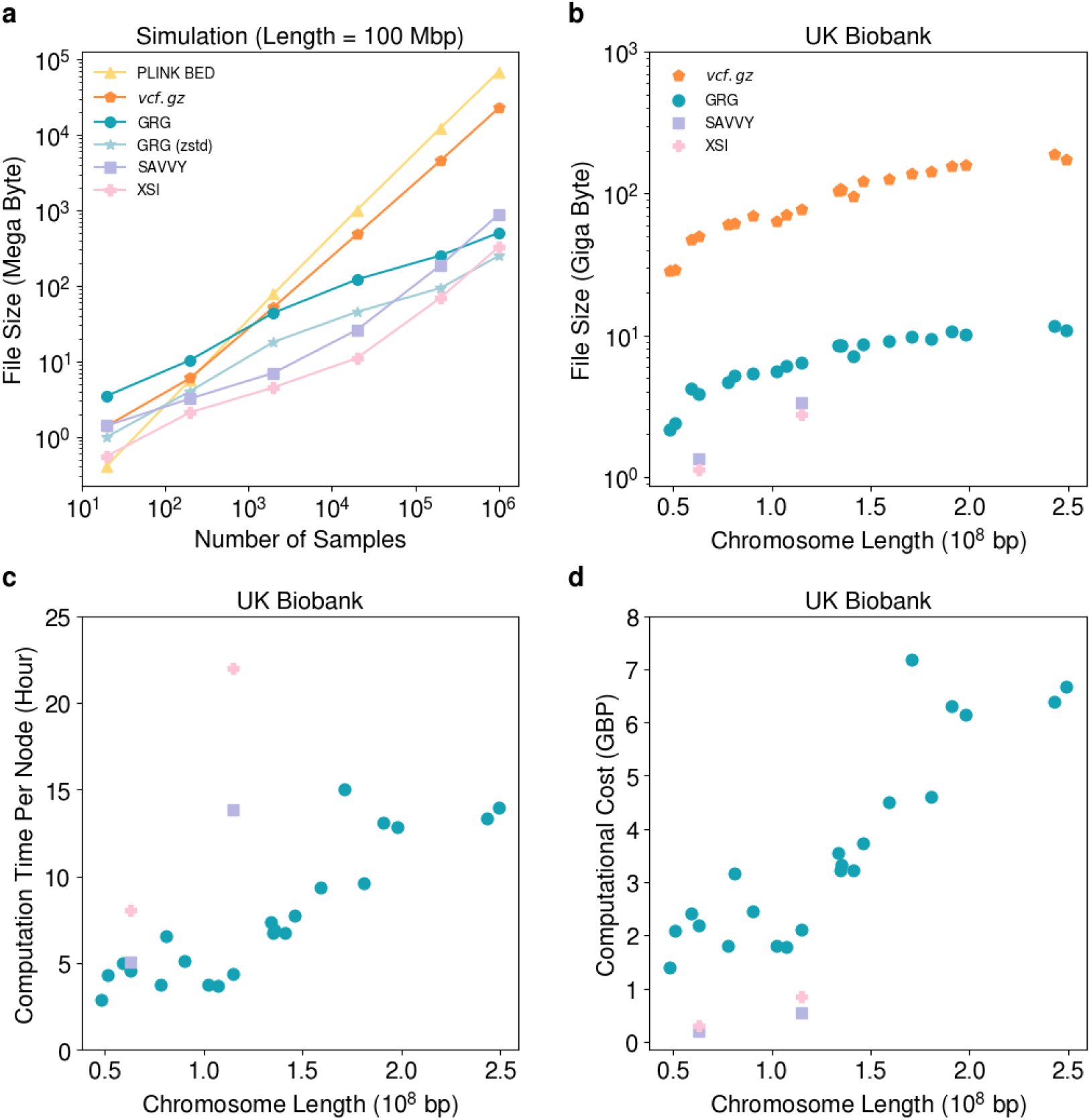
Cost, time, and file size for GRG construction. **a**: Comparisons of file sizes across PLINK BED, *vcf*.*gz*, GRG, GRG(zstd), Savvy, and XSI against different numbers of samples, with a sequence length of 10^8^ base pairs for all data points. **b**: Comparisons of GRG against *vcf*.*gz*, Savvy and XSI for storing the whole-genome polymorphisms in the UK Biobank. Each chromosome is represented by one data point. The performance of Savvy and XSI is tested with two chromosomes, 13 and 22. **c-d**: The construction time (elapsed time, not CPU time) (**c**) and computational cost (**d**) of GRG, XSI and Savvy with UK Biobank whole-genome sequencing data. The most economical cloud node was used for each method. A cloud node with up to 72 CPU threads is used for the construction of GRGs, while a cheaper node with up to 2 threads is used for constructing XSI and Savvy.

Importantly, the GRG file size scales roughly sub-linearly with respect to sample size (**Fig. 3a**). When sample sizes are smaller than 20,000, both XSI and Savvy generate files roughly half the size of uncompressed GRGs. In the dataset with 1,000,000 samples, the size of GRG is between those of XSI and Savvy, being approximately 75% larger than XSI and 33% smaller than Savvy (**Fig. 3a**). We also compared the zstd compressed GRG files, the same compression method by XSI and Savvy. With sample size 1,000,000, compressed GRG is the smallest format (**Fig. 3a**), and it is 24% smaller than XSI. However, the simplicity of having an uncompressed GRG file outweighs the small difference in file size, hence the GRG file format does not incorporate zstd or similar algorithms. The benchmarking was performed on a server with two AMD EPYC 7532 32-core CPUs (a total of 64 cores, no hyperthreading) and 1024GB of RAM.

We also applied GRG construction to all chromosomes of the UK Biobank (UKB) dataset containing 200,011 phased individual genomes <u>(Hofmeister et al., 2023)</u>. First, we converted the *vcf*.*gz* files to a fast/compact intermediate representation (see **Supplement**) and then constructed GRGs. Results are shown in **Fig. 3b-d**. GRG construction for UKB data was performed on a cloud node with 72 CPU cores and 2GB RAM per core. We used 70 of these cores during parallel GRG construction. The slowest chromosome took 14 hours to convert to GRG, resulting in a file 13 times smaller than the *vcf*.*gz*. We estimate the total uncompressed VCF size to be 350TB; large enough that decompressing it to disk is impractical. We do not filter any data; hence, the GRG represents all of the genotype information that is in the phased VCF files. The GRG files for chromosomes 1-22 store the data for more than 500 million variant sites of 200,011 phased genomes in less than 160GB. Sample identifiers are stored in the GRG file, but all other metadata for the samples and SNPs must be stored separately.

We compared the GRGs for chromosomes 13 and 22 of the UKB dataset against XSI and Savvy. Consistent with the results in simulated data, both tools produced files about half the size of the GRGs (**Fig. 3b**), with a lower file creation cost and faster construction speed (**Fig. 3cd**). Although GRG construction is approximately 15-25 fold slower than XSI and Savvy when considering CPU time, it is highly parallelizable which resulted in faster total *elapsed* construction time (using 70 threads, **Fig. 3c**) at a very reasonable cost of 80 pounds for constructing GRGs for 200,011 phased UKB genomes (**Fig. 3d**). A more detailed comparison of different file formats is provided in **Table S1** and **Fig. S4**.

### GRG as a Computational Data Structure

Dynamic programming is an algorithmic technique that exploits the storage and re-use (“memoization”) of computed values from overlapping sub-problems, and has been applied to many problems in computer science and computational biology (e.g., sequence alignment) (Giegerich, 2000; Eddy, 2004; Kelleher et al., 2016; Ralph et al., 2020; Cormen et al., 2022). The GRG graph structure resembles dynamic programming in that internal nodes represent “overlapping sub-problems”: in a compact GRG, these nodes capture the mutations-to-samples and samples-to-mutations relationship that are partially shared by many different samples and mutations. The node-sharing structure that compresses a dataset in GRG also facilitates the reuse of computed values. Computations involving multiple samples and mutations can store computed values at internal nodes, having computed those values from the graph’s parent/child edges, resulting in the same recursive overlapping sub-problems approach as dynamic programming. Dynamic programming and GRG both leverage a trade-off between storage and computation: memoization uses more storage (typically RAM) but can significantly reduce the number of computations needed.

Genome-wide summary statistics can be computed from GRG using graph traversals. The most natural traversal of a GRG is encoded in its node ID ordering: the bottom-up topological ordering of nodes in a GRG ensures that each node *n* is visited after all of its children, such that those children have node IDs less than *n*. The bottom-up ordering can also transform a recursive dynamic programming approach into an iterative approach, thus potentially accelerating computation and conserving memory in practical application.

### Allele Frequency and GWAS using GRG

To benchmark the performance of GRG-based computation, we implemented two different calculations: allele frequency and GWAS. A description of the graph traversal formulations of these calculations is provided in **Methods**, and a third graph traversal example for calculating homozygosity information is provided in the **Supplement**. We used BED files and .*vcf*.*gz* files as input to plink v.1.90b6.26 <u>(Chang et al., 2015)</u>, tree-sequence files as input to tskit (Kelleher et al., 2018), XSI (Wertenbroek et al., 2022), and Savvy (LeFaive et al., 2021), and compared the total computation time and cost against GRG (**Fig. 4a**-**c, Fig. S5**). On the simulated dataset, GRG outperformed all other data structures, including XSI and Savvy, when the sample size was larger than 200,000. For larger sample sizes, the discrepancy of efficiency between GRG and plink is expected to be more pronounced, as suggested by our simulated results. On the UKB WGS dataset, we compared plink, XSI, Savvy, and GRG on chromosomes 13 and 22 (**Fig. 4bc**). Notably, GRG achieves the fastest computation efficiency, computing allele frequencies on average more than 5-fold and 220-fold faster than plink with BED files and *vcf*.*gz* files, respectively, and more than 2.6-fold faster than XSI. A similar graph traversal approach was also used in tskit for allele frequency computations with tree sequences (Ralph et al., 2020), an ARG encoding format. Yet, GRG is more than 13 times faster on simulated data at sample size 10^6^, because “memoized” values can be reused across the entire GRG. To provide a fair comparison with the tabular formats, the GRG used here is the *constructed* GRG, whereas the tree-sequences are from the simulation (GRGs converted from simulated ARGs are even faster, see **Fig. S19**).

**Figure 4:**
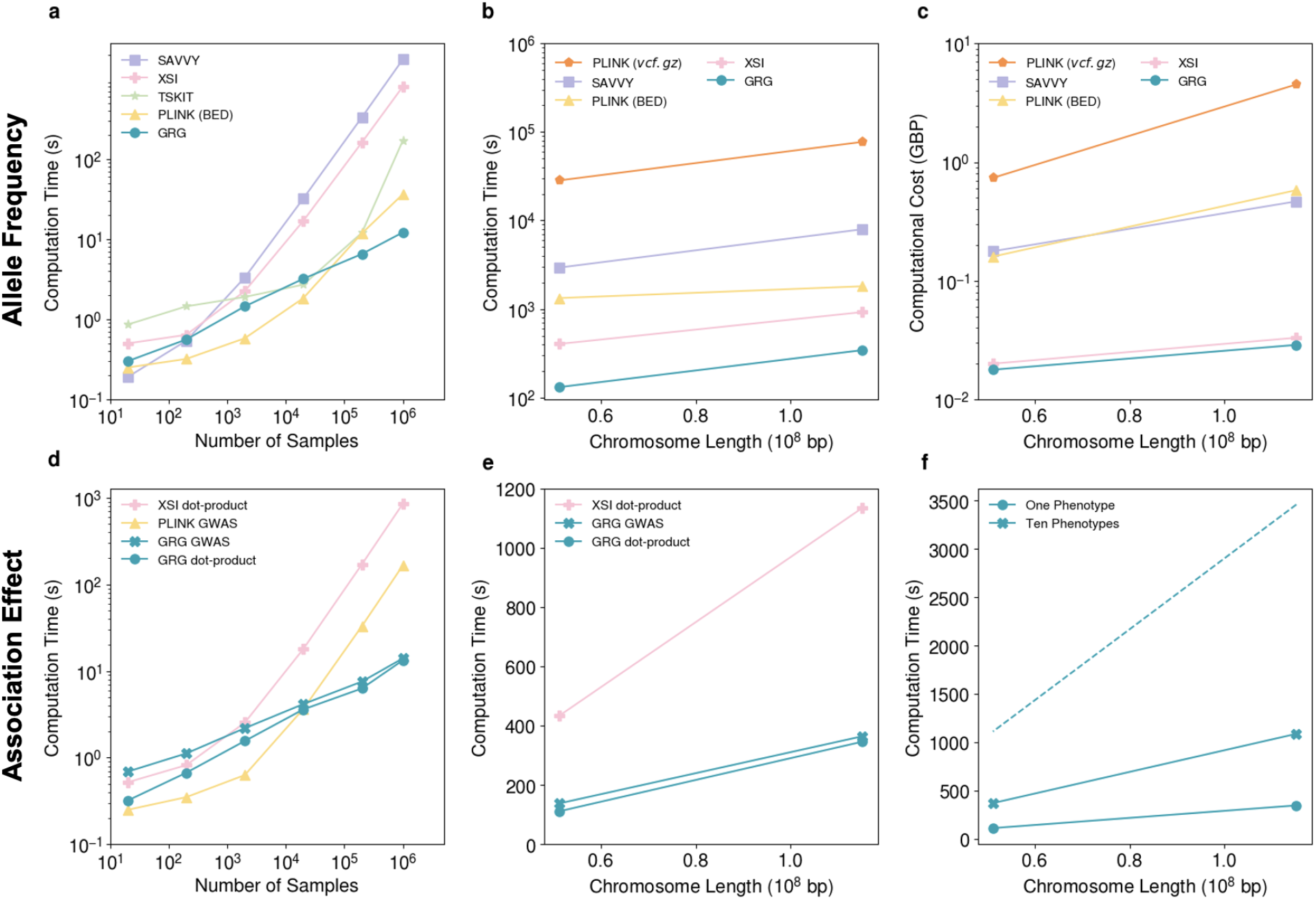
GRG accelerates genome-wide summary statistic computation. **a**: Benchmarking GRG-based computation against other methods for genome-wide allele frequencies in simulated datasets. Simulated datasets with a sequence of 10^8^ base pairs and variable sample sizes are used in this experiment. Standard tabular methods (*vcf*.*gz* and BED files) are analyzed by PLINK. XSI, Savvy and tree sequence files (tskit) are analyzed with functions from their respective libraries (see **Supplementary**). **b**-**c**: Comparison of computation time (**b**) and cloud compute cost (**c**) for allele frequencies with different data structures in the UK Biobank dataset. The most economical cloud node was used for each method. The left point of each line shows the result of chromosome 22 and the right point shows that of chromosome 13. **d**: Comparison of computation time for doing association analyses of a simulated 100 Mbp region or the dot product computation between genotypes and a phenotype vector in the simulated dataset, using matrix operations (PLINK and XSI) or graph traversals (GRG). **e**: Comparison of runtimes of the association effect computations between genotypes and a phenotype vector in the UK Biobank dataset, using XSI and GRG. Chromosomes 13 and 22 are shown. **f**: Runtimes for dot-product using GRG with varying numbers of phenotypes. Solid lines present the runtimes for loading GRGs into RAM once and running dot-product calculations with either one phenotype or ten phenotypes. The dash line is the expected time if the dot-product is calculated independently ten times.

The association effect is a measure of the genotype-phenotype relationship and is an important summary statistic in statistical genetics (Visscher et al., 2017; Abdellaoui et al., 2023). We developed a novel algorithm to compute this effect through graph traversal (see **Methods**), instead of dot product calculations via vectors or matrices. We used Genome-wide Complex Trait Analysis (GCTA) (Yang et al., 2011) to generate human-like simulated phenotypes. These phenotypic values, together with genotypes encoded in the respective data structures, were used for genome-wide association studies. We provide two modes of calculation: full association results with statistical testing, or only the association effect sizes (which we refer to as “dot product”). The former is for comparison with PLINK and the latter for comparison with XSI (which provides a comparable dot-product implementation) (**Fig. 4df**). We verified that our graph traversal approach yields association effect, standard error, and p-value identical to the commonly used software plink <u>(Purcell et al., 2007)</u>. In the simulated dataset, our results indicate that when sample sizes are larger than 20,000, the graph traversal algorithm consistently outperforms the matrix operations in terms of speed, regardless of whether the data is encoded in a traditional tabular format or a compressed sparse format. On the UKB dataset, GRG performs the dot-product calculations more than 3 times faster than XSI, even though the GRG file is more than twice the size of the XSI file. The difference in computation time between GRG and XSI/Savvy is more prominent in the simulated data than in the UKB data, which could be due to a combination of the computer performance (UKB experiments were done in the cloud) and differences in the variant distribution between the two datasets (UKB is significantly denser with variants, and in particular has a higher proportion of low-frequency alleles). All computation times in our figures *include* the time to load the GRG from disk, but **Fig. 4f** demonstrates that performing multiple computations after loading the GRG into RAM is significantly faster than loading from disk each time.

### Biological Explanations of the GRG Multitree Data Structure

The GRG data structure belongs to a data structure group that is often referred to as a multitree or strongly unambiguous graph, where there is at most one direct path between any two nodes (Furnas and Zacks, 1994). A unique feature of the multitree data structure is that each node in the graph has a tree both above and below it, making it particularly suitable for leveraging any tree-based algorithms. Although GRG construction is motivated by its computational benefits, we are also interested in the potential biological interpretations of the GRG data structure. From a coalescent theory perspective, it is intuitive that a descendant tree structure is a natural way to encode inheritance of variants from a common ancestor to present-day samples (Wakeley, 2009). To our knowledge, the existence of a parent tree structure has not been discussed in the coalescent theory literature.

Different from the coalescent with recombination model (Hudson, 1983a; Hudson, 1983b) that focuses on modeling each evolutionary event (i.e., mutation, recombination, coalescence), the multitree structure in a GRG can be seen as a natural encoding of newly arised haplotypes. As previously mentioned, each GRG node represents a haplotype consisting of all the mutations in its parent tree. In the forward-in-time evolutionary process that generates present day sampled genomes, many new haplotypes are created over time, either due to new mutations or recombinations. Here, we define a GRG haplotype as a unique combination of mutations that are co-inherited by one or more present-day samples through a unique (coalescent) tree. By this definition, a GRG haplotype could be one or more mutations contained by a GRG root node, a combination of mutations reachable from a GRG internal node upward (inclusive), or a full length present-day chromosome reachable from a GRG sample node (**Fig. 5a**). In **Fig. 5a**, we illustrate a hypothetical pedigree history of a focal GRG haplotype. If we trace only the genetic materials in this focal haplotype back to its genetic ancestors, a tree above this haplotype arises. This tree represents how genetic materials are transmitted to the focal haplotype during evolution. Any new mutations that occur during transmission are also represented by the same tree. Therefore, the parent tree structure of a GRG node could be explained based on the formation of a new haplotype. The descendant structure of a GRG node is also a tree – consider a unique GRG haplotype being transmitted to some present day samples, these samples are related to each other through a coalescent tree.

**Figure 5:**
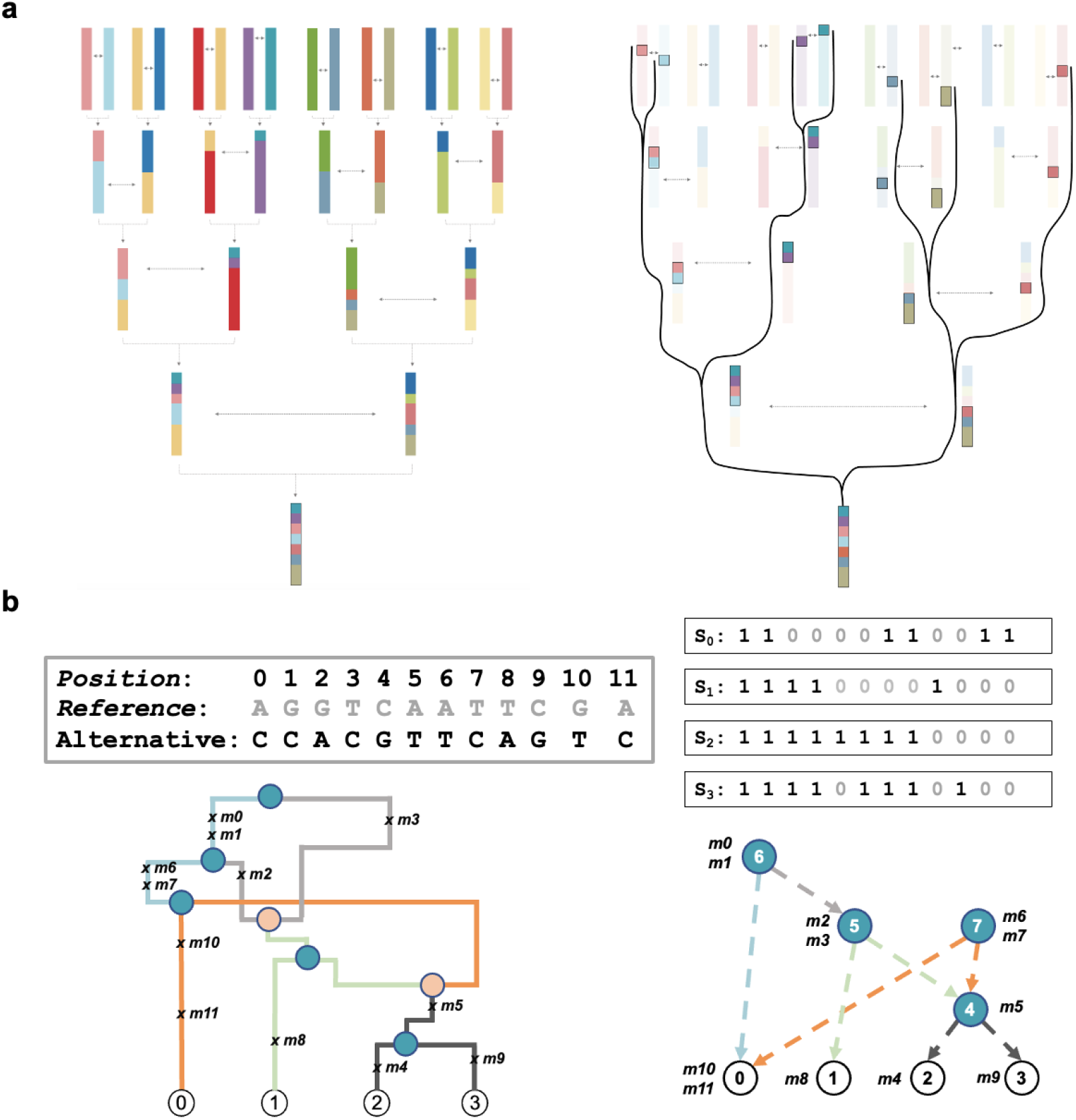
The biological interpretation of the GRG data structure. **a**: The tree above a GRG node can originate from the pedigree inheritance. The left panel shows the full pedigree of a focal haplotype, including the ancestral inheritance and recombination events. The right panel shows the specific haplotypes in the pedigree that are passed down to the focal haplotype, and the paths to the focal haplotype form a tree shape. Here, recombination events cause the merging of lineages. **b**: GRG as a reduction of ARG. The reference and sample haplotypes are listed. Here, we highlight the genealogies in ARG that remain in the corresponding GRG. When present in an ARG, the following redundant structures are removed to become a GRG: 1) edges and nodes that represent the same topological structure with different coalescence time (e.g., one of the two light gray edges in the ARG is removed); 2) edges without mutations (e.g., the upper green edge in the ARG is removed); 3) mutations that are inherited by the same set of haplotypes through different paths are merged (e.g., *m2* and *m3*).

Depending on how it is constructed, a GRG can be seen as the graph that encodes how new genetic information is created in the evolutionary process that generates sampled genomes. Similar to a phylogenetic tree, how well a constructed GRG actually captures this evolutionary process depends on the construction algorithm. In this work we have developed a greedy algorithm akin to parsimony in phylogenetic tree construction, but going forward we have interest in constructing better GRGs to capture the haplotype evolution process.

### ARG to GRG Conversion

In this work, we focus on scalable GRG construction to facilitate data compression and scalable computation, rather than inferring the actual haplotype formation process during evolution; hence, we refer to our method as GRG construction rather than GRG inference. Because a GRG encodes less evolutionary information than an ARG, and ARG inference is aimed at capturing the true genealogical history, we can also convert an ARG into a GRG that can better resemble the haplotype formation process (**Fig. 5b**). It is important not to confuse GRG being a reduction from ARG with GRG being a reduction from tree sequence (TS), which is a popular ARG *representation* <u>(Kelleher et al., 2019)</u>. In practice, tree sequences can be converted to GRGs (see **Supplement** for the algorithm).

Conversion from TS to GRG is extremely fast (about 30 seconds for 1 million haploid samples of length 100Mbp, see **Fig. S6**), and could be useful when the dataset is quite large or computations will be performed many times. As the number of samples gets really large, GRG can perform computations like allele frequency much faster than TS (**Fig. 4a**), because a value memoized at a specific node is usable across the entire genome, whereas with TS memoized values may only be valid for a sub-region of the genome. Another potential benefit of TS to GRG conversion is to speed up analyses (e.g., **Fig. 4**) that simply rely on the haplotype history among samples (but not any additional information on an ARG), as such information could be useful in population genetics.

Because a GRG could either be directly constructed from a genotype matrix or be converted from an ARG, we evaluated which GRG can compress the data better (**Fig. S7**). In simulated data, the constructed GRG is larger than the TS-GRG (**Fig S7ab)**, but the opposite is true in the 1000 Genomes Project data (**Fig S7cd**). Our additional experiments suggest that this discrepancy could be due to errors in the real data influencing the robustness of ARG reconstruction and GRG construction disproportionately (**Fig. S8**). When a GRG is converted from an ARG (**Fig. S7ab**), the size of the GRG increases sublinearly with sample size, whereas the trend is a bit less obvious with constructed GRGs (**Fig. S7ab**., **Fig. 3a**).

### Discussion

We presented a new data structure and file format termed genotype representation graph (GRG) and an algorithm for GRG construction. Constructing GRGs can be done for biobank-scale datasets in a matter of days. GRG construction is much faster and more scalable than ARG inference (**Fig. S7d**), though slower than PBWT-based compression approaches. We showed that calculations can be performed much more efficiently via GRG than other file formats, for biobank-scale datasets. This speed for calculation also makes GRG a useful representation for simulated population genetic data, especially since converting from the tree sequence format to the GRG format is extremely fast - on the order of a few minutes for very large datasets. We also presented a graph-based algorithm for doing genome-wide association studies (GWAS). To our knowledge, this is the first graph-based GWAS where each diploid individual has a phenotypic value (see (Ralph et al., 2020)) for a simpler approach that assumes each haploid sample has a phenotypic value), practical for analyzing large-scale human datasets.

Compared to other file formats (Purcell et al., 2007; Durbin, 2014) such as BED files which prohibit multi-allelic polymorphisms, GRG provides a unified way of presenting both bi-allelic and multi-allelic polymorphisms as mutation nodes. Similarly, GRG can encode insertions and deletions with mutation nodes. Missing data can be handled by GRG as separate mutation nodes that are marked specially. Whereas a typical mutation is a pair (genomic position, alternate allele), a missing data mutation is a pair (genomic position, *missing data flag*). When there is a lot of random missingness in the dataset, the graph size can increase non-trivially, since it needs to encode these additional mutation-to-sample relationships. In the UK Biobank SHAPEIT phased whole genomes (Ribeiro et al., 2023), there is no missingness, as missing genotype imputation is usually coupled into the phasing pipeline (Howie et al., 2009; Hofmeister et al., 2023).

Generally speaking, the purpose of the GRG data structure is to compactly and faithfully encode haplotypes as mutations being passed down via paths in the graph, to allow easy access to both mutation information and haplotype information. In contrast, ARG data structures are designed to fully record the true or inferred inheritance history of the genomes, including coalescence times (Hudson, 1990; Griffiths and Marjoram, 1996; Griffiths and Marjoram, 1997; Rasmussen et al., 2014; Kelleher et al., 2016). GRG supports random queries of variant-to-sample relationships and sample-to-variant relationships – querying a variant-to-sample relationship simply requires traversing downward from the node bearing the variant to collect sample nodes, and querying a sample haplotype simply requires traversing upward from the sample node to collect mutations (**Fig. S9**). Simple tabular data formats allow easy access to either variant-to-sample relationships or sample-to-variant relationships but not both. Indexed tabular formats such as XSI and Savvy do provide the variant-to-sample relationship, and perform calculations on small genomic regions (less than 1Mbp) faster than GRG (**Fig. S10**). For larger genomic regions or subsets of samples, GRG is faster (**Fig. S10**), therefore, particularly suitable for large-scale analyses. Unlike other formats, GRG can usefully store massive datasets in RAM because the graph representation *is the compression*, meaning there is no decompression step. Repeated computations or interactive data analysis (such as via Jupyter Notebook) are then significantly faster than other compressed formats, as graph traversal times can be orders of magnitude smaller than the time to load GRG from disk (**Fig. 4f, Fig**.

**S10**).

Coincidentally to what we have observed with the UK Biobank data (**Fig. 3**), in astronomy, multitree methods typically yield orders of magnitude in speedup over the alternatives, making millions of data points tractable on desktop workstations (Gray et al., 2004). We, therefore, anticipate that the GRG data structure can also broadly improve the scalability of data analyses involving large genetic datasets. The GRG dot product (**Fig. 4de**) provides an essential building block for doing more complicated statistical genetic work. To facilitate the adoption of GRG in other studies, our library’s API provides visitor and iteration patterns for depth-first, breadth-first, and topological order traversals. The API is available in Python for ease-of-use, and C++ for users needing higher performance.

Both GRG construction and ARG inference can generate a graph that encodes all the information in a genotype matrix. For representing the true ARG, the tree sequence format is currently the most compact format. In terms of compressing genotype matrices and facilitating genotype analyses, GRG can be superior as GRG construction is more scalable (**Fig. S7**) and it is more robust to errors in real data (**Fig. S8**). In real datasets, sequencing and genotyping errors are orders of magnitude larger than mutation rates, with many error rate estimates near 0.1% (Saunders et al., 2007; Browning and Yu, 2009; Halldorsson et al., 2022) (see **Supplement**). We suspect that there is room for further reducing GRG sizes by generally improving robustness to noise, or by identifying and “ignoring” errors. In general, sequencing errors appear to pose a challenge for any technique that is trying to identify the true inheritance relationships between haplotypes, whether for the purpose of compression (GRG) or genealogy inference (ARG).

The time to construct a GRG and the GRG file size both roughly follow the number of polymorphic sites in a dataset, even when the data comes from populations with different ancestry (see **Fig. S11**). Notably, genotyping array data and whole-exome sequencing data are still frequently used. These two types of data are designed to measure only a small fraction of the genetic polymorphisms, hence are computationally inexpensive compared to whole-genome sequencing data. We expect GRG to be able to compress these data as well, but to a lesser extent.

Although we primarily use the UK Biobank data as an example, the utility of GRG is not limited to human data. Indeed, data from other species are also rapidly growing. We converted two sets of inferred tree sequences for the SARS-CoV-2 genomes (Zhan et al., 2023) into GRGs, one for 1.27 million SARS-CoV-2 genomes sampled up to June 30, 2021, and the other for sparsely sampled 657 thousand samples sampled up to June 30, 2022, both originally from GISAID (Shu and McCauley, 2017) (**Table S2**). For these two tree sequences the resulting size of the GRG is similar to the size of tree sequence format (after discounting meta-data).

In this work, we focus on GRG construction and its potential utility in compression and GWAS, but we also explore a connection between the GRG data structure and the genetic data generating process. The haplotype view of the GRG data structure is interesting because it could be useful in population genetic analyses. Evaluating the evolutionary properties of a GRG constructed by a particular algorithm, and exploring the space of GRG construction algorithms could be an interesting and challenging future direction. Because our current GRG construction algorithm resembles ARG inference and evolutionary tree building, another future direction could be to investigate whether GRGs can help facilitate more scalable ARG inference.

Generative models in machine learning have demonstrated exceptional performance in tasks like image synthesis, text generation, and data augmentation (Gm et al., 2020). Extending their utility, these models can be applied to lossy data compression by generating compressed data representations (Kingma et al., 2019; Yang et al., 2022). However, such compressed representations cannot support the same computation as with uncompressed data. In population genetics, researchers have long employed models motivated by the mechanisms in the biological world to closely resemble the true genetic data-generating process to gain insights into genetic polymorphism. The construction of a GRG can be seen as an approximation to this data-generating process to losslessly compress the observed genetic polymorphism. In addition to achieving a highly compressed data representation, GRG improves the ability to support computations on the original data. Our data structure and algorithm, inspired by population genetics principles, addresses computationally challenging problems beyond the traditional scope of population genetics.

### Methods

#### BuildShape

Given a dataset, let *S* = {0*…N-*1} be the set of all samples and *M* = {*M*_0_*…M*_V_} be the set of all mutations. Let *S*_i_ refer to the subset of samples from *S* that contain the *i*-th mutation, *M*_i_. We’ll use 𝒢 to refer to a corresponding GRG. The GRG construction problem is to search from the space of all possible internal nodes to find an optimal set of internal nodes such that 1) all mutations in *M* are correctly mapped to a node and 2) the total number of edges in 𝒢 is minimized.

*BuildShape* constructs an initial graph 𝒢 by iterating over a set of nodes *S’*. Initially *S’* = *S*, i.e. the set of all sample nodes. For each sample *i*, we find the sample *j* with the most similar haplotype. We create the union of these samples by constructing a new node *N*_k_ and making it the parent of nodes *i* and *j*. Nodes *i* and *j* are removed from *S’*, and node *N*_k_ is added (i.e., *S’* = *S’* \ {*i, j*} ∪ {*k*}). The order of samples could influence the final graph.

Building the initial GRG shape requires *O*(*N*^2^) haplotype comparisons, where *N* is the number of samples. We speed up the Hamming distance calculation and reduce memory footprint by reducing the dimensionality of each haplotype. Instead of exactly representing mutation lists, we use bloom filters (Bloom, 1970) with a single hash function (*k* = 1) and bit-size *m* = *A* × *B* where *A* is the average number of mutations per sample and *B* is a parameter specifying how many bits to use for each mutation (see **Supplement**). We also use a BK-tree (Burkhard and Keller, 1973) nearest-neighbor index to speed up the similarity search.

The *T* GRGs that cover the genome are merged into *P* <= *T* sub-GRGs prior to *MapMutations*. For simplicity, **Fig. 2a** shows the case where *P* = *T. P* determines how parallelizable GRG construction is: each merged sub-GRG can have mutations in the corresponding genomic region mapped onto it in parallel. In practice, *P* is expected to be larger than the number of cores available for GRG construction. An increase in *P* only marginally increases the size of the final GRG, and larger *P* can reduce overall runtime even when construction is run on a single core (see **Fig. S3cd**).

#### Determining Parameters T and P

*T* is the number of GRG-trees to construct for the genome, an important hyperparameter that determines the construction efficiency. Varying this parameter can significantly affect graph size (**Fig. S3ab**). If *T* is too large (segment size is small), there may not be enough variation in the distribution of mutations in that region to accurately ascertain the genealogy among samples. If *T* is too small (segment size is large), the important sample relationships cannot be well-described by a single tree, due to recombination within the region. We split the genome evenly, according to base pairs, between the *T* trees. We’ve found that for real data the optimal segment length is in the range 50 - 150Kbp, which means that the optimal *T* varies depending on chromosome length. The lower range (50Kbp) works best for data that is dense in mutations-per-sample (such as the 1000 Genomes Project). The upper range (150Kbp) works better for sample-rich data (such as the UKB). A method for finding the best parameter for a new dataset is to simply construct multiple GRGs for a small 500Kbp segment of the genome: one for each segment length in the set {50, 75, 100, 125, 150}. Once a reasonable segment length for a particular chromosome in a dataset is found, it can usually be reused across all chromosomes.

In our experiments, simulated data performs better with much smaller *T* values than non-simulated data. Adding noise to the simulated data did not have a significant impact on the optimal *T* value, so it is not clear what causes this difference between real and simulated data.

In contrast to *T*, the *P* parameter has very little effect on the final graph size, but has a significant effect on runtime (**Fig. S3**). *P* must be less than or equal to *T* and should be greater than or equal to the number of cores available for GRG construction.

#### MapMutations

*MapMutations* modifies each sub-GRG such that each *M*_i_ in *M* is assigned to a node that covers exactly *S*_i_. In the worst case, a new node *N*_k_ is added for every *M*_i_, and edges are added from *N*_k_ to every sample node in *S*_i_. For each mutation *M*_i_, MapMutations does an upward graph search starting at *S*_i_ and collects all nodes that have *only* samples in *S*_i_ beneath them. In the best case, it finds an existing node that exactly covers *S*_i_ and just assigns *M*_i_ to that node. More typically, a new node is created for the mutation and multiple nodes *N*_j_ are discovered such that the union of samples covered by *N*_j_ is a subset of *S*_i_ and the intersection of samples covered by *N*_j_ is empty. The empty intersection ensures we maintain GRG property **V** (no diamond patterns).

By default, when mapping mutations we calculate the frequency of each alternative allele and swap the reference allele to the major allele when an alternative allele exceeds frequency 0.5. This allele flipping behavior can be disabled if it is preferred to keep the original reference, for example when the dataset has been polarized.

#### Merging GRGs

When two GRGs are merged the result is simply the union of the nodes and edges, except that internal nodes containing the *exact same* covered sample set are combined into a single node. We perform this merge by mapping the second GRG onto the first. The first GRG is traversed and a one-way hash is computed of the samples covered by each node. Then the second GRG is traversed in bottom-up topological order and the same hash is computed for each node. Each node and its edges are copied to the first GRG *unless* the node has a matching one-way hash in the first GRG. Regardless of whether the hashes match, the mutations are always copied from the second GRG to the first.

Node renumbering, graph simplification, Compressed Sparse Row (CSR) encoding (Duff, 1984; Buluç et al. 2009) (**Fig. S12**), and serialization to disk are then performed simultaneously. A depth-first search, using arbitrary edge ordering, is performed over the entire graph to produce a topological ordering such that the sample nodes take the node orders 0…*n-*1 and all other nodes are ordered behind them. The nodes are iterated in this order, and a *node position vector* and an *edge vector* are constructed. The node position vector value at index *i+*1 holds the index *e*_i_ in the edge vector that contains the first edge for node *i*. If node *i* has *c* down edges, then positions *e*_i_…(*e*_i_ + *c*) of the edge vector contain those edges, where each edge is represented by the node ID *j* for the edge (*i, j*).

GRG is both an in-memory data structure and a binary file format. The node ordering and CSR encoding mean that data for all nodes and edges are stored sequentially (**Fig. S12**), such that traversing the graph from the samples to the mutations can be done easily (simply iterating over the node identifiers) and cache-efficiently. Using this custom format we can store huge datasets much more compactly and efficiently than using a standard graph file format like GraphML (Brandes et al., 2002).

#### Simulating Ground Truth Tree Sequences

To investigate the performance of different data encodings, we simulated population genetic polymorphisms under different conditions. Msprime v.1.2.0 <u>(Baumdicker et al., 2022)</u> is used to run a series of backward-time coalescent simulations to generate the ARGs encoded as tree sequences (Kelleher et al., 2016; Kelleher et al., 2018) with varied (diploid) sample sizes ranging from 10 to 500,000. For simulated data presented in the main text, we set a constant effective population size (Ne = 10^4^), a flat recombination rate of 10^−8^ / (bp × generation), and a mutation rate of 10^−8^ / (bp × generation). The random seed is 42 for all simulations. More complicated models including human demography, gene conversion, and variable recombination rate are also tested (see **Supplement**).

We convert the tree sequences into phased VCF files with tskit v.0.5.5 (Kelleher et al., 2018), and convert the phased VCF files into bed files with plink v.1.90b6.26 (Chang et al., 2015).

#### Constructing GRGs from non-Simulated Data

We used two datasets for GRG construction: the 1000 Genomes Project (The 1000 Genomes Project Consortium, 2015) and the UKB data (Hofmeister et al., 2023).

By default, our 1000 Genomes Project experiments use all 2504 individuals from the Phase III data (The 1000 Genomes Project Consortium, 2015), unless stated otherwise. We used parameters *T* = 1600 and *P* = 50 for GRG construction. All of our experiments used 20 CPU threads during GRG construction.

All UKB GRG construction was performed in the cloud using the Research Analysis Platform, and the SHAPEIT phased VCFs from the “GATK and GraphTyper WGS” release containing genotypes for 200,011 individuals (McKenna et al., 2010; Eggertsson et al., 2017; Ribeiro et al., 2023). We used 70 CPU threads during GRG construction for chromosomes 1-22. We used various *T* values from 280 (chromosome 22) to 1050 (chromosome 1), and *P* from 70 (chromosome 22) to 350 (chromosome 1).

#### Allele Frequency Computation

For tabular data, allele frequency computation can be done element by element (**Fig. 6a**). In a GRG, the sample frequency at any node *n* can be computed as the sum of the sample frequencies for each child node (**Fig. 6b**). Since the children of node *n* have IDs less than *n*, we can simply compute sample frequencies in the order of nodes 0…*n* and store each frequency. The frequency values of all mutation nodes can then be extracted to obtain the allele frequencies for the dataset. This process guarantees that mutations containing the same subset of samples (*S*_1_, …, *S*_k_) captured by an internal node will all reuse this internal node’s frequency.

**Figure 6:**
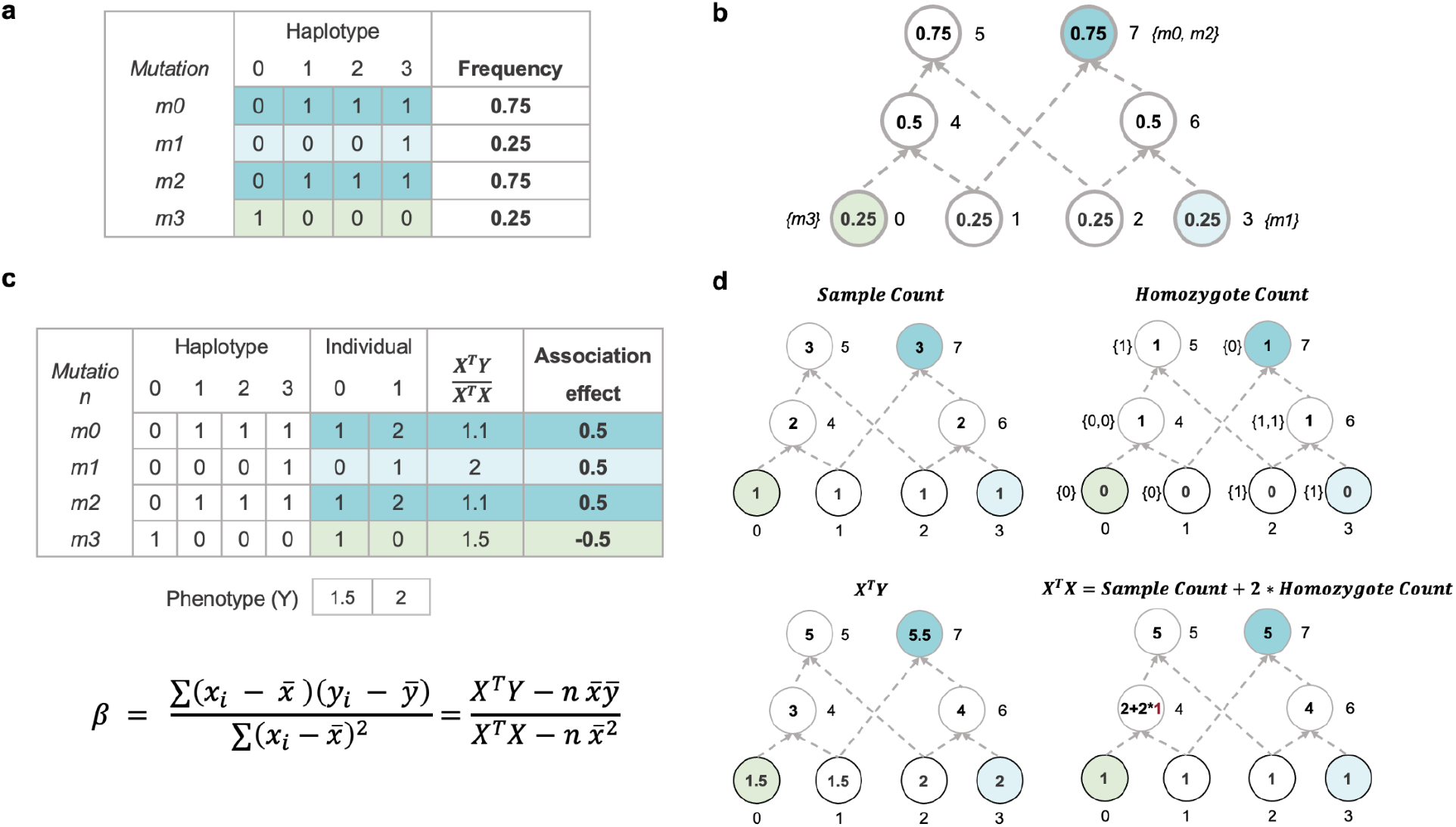
Schematic illustrations of graph-based algorithms for computing allele frequencies and association effects. **a**: Computation of allele frequencies using standard tabular methods. Genotypes of each variant are read row by row from disk and allele frequencies are computed independently for each variant in the tabular structure. **b**: Computation of allele frequencies using GRG. In a GRG, the sample frequencies of nodes are firstly computed and then mapped to the mutations based on the node-mutation mapping. The frequency of leaf nodes is set as 1/n (e.g., node0-3), where n is the number of haplotypes in the population. For other nodes (e.g., internal nodes and root nodes), frequencies are computed sequentially based on the node IDs (e.g., from node4 to node7) as the sum of frequencies of each node’s children nodes (e.g., node4 = node0 + node1). **c**: Computing association effect (β) with genotype and phenotype vectors. **d**: A schematic illustration of computing association effect through graph traversal. The *sample count* is the count of haplotypes beneath a node. The *homozygote count* can be propagated using the *GRG*.*numIndividualCoals* via graph traversal. The computation of *X*^*T*^Y is similar to the sample count, except that the initialized values for leaf nodes (labeled inside the leaf nodes) are the phenotypic values of the sample nodes at the individual level. The term *X*^*T*^*X* is computed as *sample count* + 2 × *homozygote count*.

#### GWAS Computation

The full equation is given by:

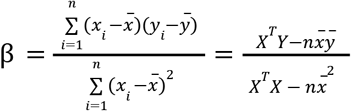

Here, *X* is the vector of genotypes of *n* diploid individuals at a mutation, *Y* is the vector of phenotypic values of individuals. The *X* vector has a single value for each individual indicating the number of copies of the alternative allele that are present (0, 1, or 2).

The term *X*^*T*^*X* can be decomposed further into the counts of homozygotes 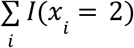 and the counts of heterozygotes 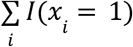, where I is the indicator function. The ancestral allele homozygotes *x*_*i*_ = 0 terms are not present in the GRG, and do not contribute to the dot product result. Each homozygote contributes a value of 4 to the *X*^*T*^*X* term, while each heterozygote contributes 1. Therefore, we can derive:

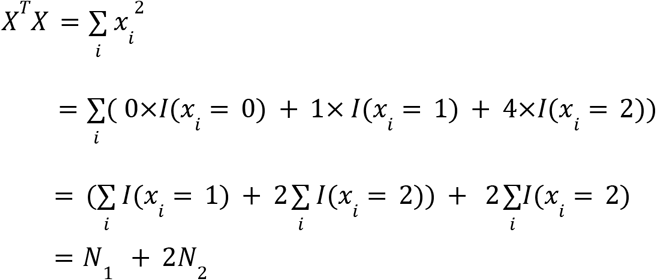

*N*_1_ and *N*_2_ can be interpreted in the graph as the number of samples and the number of homozygous individuals reachable from a node. Therefore, the equation can be further revised as:

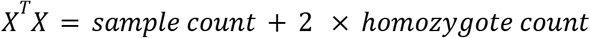

The GRG itself stores *numIndividualCoals* at each node, which is the number of individuals that coalesce *exactly* at that node. By propagating the values *numIndividualCoals* of all children to their parent node (**Fig. 6c**), our algorithm produces the *Homozygote Count* for the derived/alternative allele as obtained from the genotype matrices (also see **Supplement**).

All additional terms can be easily computed or accessed. The *X*^*T*^*Y* term computes the dot product of individuals’ genotypes and phenotypes. Similar to allele frequency computation, the *X*^*T*^*Y* term can be computed by a single graph traversal, where sample nodes are initialized with their respective phenotypic values at the individual level. Similarly, the average of *X* vector, 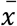, can be computed via graph traversal. The average phenotype 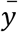 is computed when loading the phenotypes. The term *n* is given in a GRG.

The GRG-GWAS computation yields two modes of results: association effect only or full association testing. In addition to the association effect β, full association testing generates β_0_, standard error, *R*^2^, t-value, and p-value. All of these can be easily computed based on the aforementioned terms (*n*, 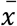, 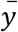, *X*^*T*^*Y*, and *X*^*T*^*X*) used for computing β, in addition to *Y*^*T*^*Y* which can be computed when loading the phenotypes. β_0_ is computed through the equation:

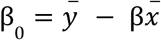

The sum of squares residual error term (*sse*) of linear regression is

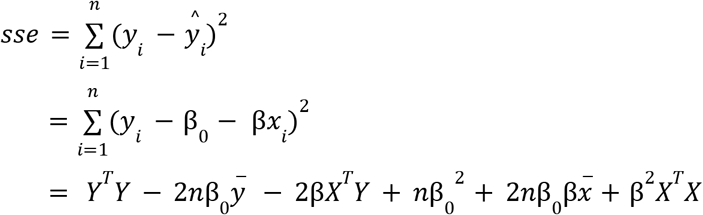

The sum of squares total is:

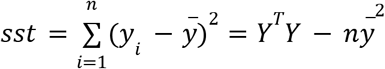

The *R*^2^ for the regression is:

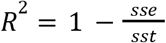

The standard error of association effect is:

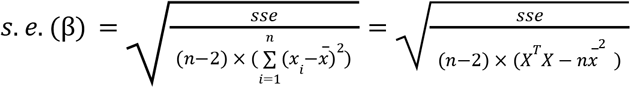

The t-value of association effect is:

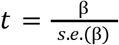

The p-value of association effect is computed using the GNU Scientific Library (GSL) (Galassi 2009) with the *gsl_cdf_tdist_P* function. The t-value and degree of freedom equal to n-2 are used as input to the *gsl_cdf_tdist_P* function.

#### Inferring Tree Sequences from Genotype Matrices

To infer tree sequences from genotype matrices or true tree sequences, we use tsinfer v.0.2.3 (Kelleher et al., 2019), a scalable tree inference method. True tree sequences can be easily converted to the .*sample* data format required by *tsinfer* using the command *tsinfer*.*SampleDate*.*from_tree_sequence* in the python API. To convert genotype matrices (phased VCF files) to .*sample* files, a python script is implemented to load the genetic variants in VCF files with the cyvcf2 package (Pedersen and Quinlan, 2017) and convert them into the sample object. Per the guideline of *tsinfer*, only biallelic sites can be used for inference. For positions with multiple allelic variants, only the reference allele and the first alternative allele are retained in the .*sample* files. Non-single nucleotide polymorphisms are also removed. Once the .*sample* files are prepared, the *tsinfer*.*infer* function is used to estimate the genealogies with the full inference pipeline.Default settings are applied for the recombination rate and mismatch ratio.

For the 1000 Genomes Project, we used 20 threads during *tsinfer* inference for chromosomes 1-3, 10, 13-15, and 21; 10 threads for chromosomes 4-9, 11-12, and 22; and 5 threads for chromosomes 16-20. For the simulated data, we used 5 threads for simulations with sample sizes from 10 to 10,000, and 10 threads for 100,000 samples.

#### Tree Sequence to GRG Conversion

Tree sequences are converted to GRG by copying and duplicating nodes and edges from the tree sequence into a new GRG. Every recombination event that occurs between trees 𝒯_k_ and 𝒯_k+1_ in a tree sequence can be represented as a set of edge insertions and deletions to 𝒯_k_ in order to produce 𝒯_k+1_. Consider a single edge *E*_1_ = (*n*_i_, *n*_j_) being deleted and another edge *E*_2_ = (*n*_r_, *n*_p_) being added (**Fig. S13**). *n*_i_ and *n*_r_ are the parent nodes for these edges. The nodes above *n*_r_ now potentially have more samples beneath them (because *n*_p_ is a new child, and it has samples beneath it). The nodes above *n*_i_ now potentially have *fewer* samples beneath them (because *n*_j_ is no longer a child, and it had samples beneath it). Now consider the most recent common ancestor (MRCA) (Kingman, 1982b; 1982a) between *n*_i_ and *n*_r_. All nodes at and above the MRCA have no change to their reached samples: those samples were moved from one subtree of the MRCA to another subtree of the MRCA. This implies an efficient tree sequence to GRG conversion algorithm: nodes from 𝒯_0_ are copied into GRG 𝒢 to produce the initial graph. The MRCA between added/removed edges for subsequent trees 𝒯_i_ is used to determine a list of nodes that need their reached samples modified. In constructing the GRG, we duplicate a node whenever its reached sample set changes, and copy the relevant graph edges.

Even for large inferred tree sequences, converting from a tree sequence to a GRG takes on the order of minutes. Converting simulated tree sequences to GRG takes seconds even for hundreds of thousands of samples.

## Data Availability

The phased *vcf*.*gz* files used in this study from the Phase III 1000 Genomes Project can be downloaded through https://hgdownload.cse.ucsc.edu/gbdb/hg19/1000Genomes/phase3/.

The UK Biobank data is available to approved researchers through http://dnanexus.com/. The latest SHAPEIT phased VCFs from the field 20279 are used for analysis and the details about it are documented at https://biobank.ndph.ox.ac.uk/showcase/refer.cgi?id=1910. The constructed GRG from the SHAPEIT phased data will be returned to the UK Biobank for access by approved projects.

The inferred tree sequences of SARS-CoV-2 genomes from Zhan et al., 2023 are available by request from the original authors with approved GISAID data access. The original genomes of the SARS-CoV-2 are available on GISAID, via gisaid.org/EPI_SET_230329cd and included in the Supplemental Table (GISAID EPI SET PDF). The converted GRG of SARS-CoV-2 genomes are available by request with approved GISAID data access.

## Code Availability

We have released an open-source, lightweight VCF parser picovcf v2.0 through https://github.com/aprilweilab/picovcf/releases/tag/v2.0. The Genotype Representation Graph Library (GRGL), which implements the construction of GRGs and downstream computations, is available through https://github.com/aprilweilab/grgl. A detailed description of the software is provided in the **Supplement**.

External softwares used in this study include:

msprime v.1.2.0, https://pypi.org/project/msprime/1.2.0/

tskit v.0.5.5, https://github.com/tskit-dev/tskit/releases/tag/0.5.5

tsinfer v.0.2.3, https://github.com/tskit-dev/tsinfer/releases/tag/0.2.3

plink v.1.90b6.26, https://www.cog-genomics.org/plink/.

XSI v4.0.1 (dev): https://github.com/rwk-unil/xSqueezeIt/tree/21562519

Savvy v2.1.0: https://github.com/statgen/savvy/releases/tag/v2.1.0

GCTA v.1.94.1: https://yanglab.westlake.edu.cn/software/gcta/bin/gcta-1.94.1-linux-kernel-3-x86_64.zip

## Supplementary Materials

Supplementary Text, Algorithm S1-S4, Table S1-S2, and Figure S1-S19, and a Supplementary Table for GISAID SARS-CoV-2 data.

## Acknowledgments

We gratefully acknowledge all data contributors, i.e., the Authors and their Originating laboratories responsible for obtaining the specimens, and their Submitting laboratories for generating the genetic sequence and metadata and sharing via the GISAID Initiative, on which this research is based. We thank Jerome Kelleher for generously sharing the inferred SARS-CoV-2 tree sequences with us. This work is partly supported by NIH R35-GM150579 to X.W. This research is conducted under the UK Biobank application 97908.

