## Supplement for "Genotype Representation Graphs: Enabling Efficient Analysis of Biobank-Scale Data"

### Genotype Representation Graph Library (GRGL)

Our implementation of GRG is distributed as a library that can be used either from C++ or Python code. The library also includes command-line tools for converting other file formats to GRG and computing simple statistics for a GRG.

All nodes are represented by a unique *NodeID*, an integer between 0 - ($n$-1), where $n$ is the number of nodes. All node-based operations take a *NodeID* as an argument. There are two different modes of graph traversal:

1. Non-terminating graph traversals can be done by iterating *NodeID*s. Just iterating from 0 … ($n$-1) will traverse the entire graph in bottom-up topological order (equivalent to performing a depth-first search that uses an arbitrary child order at each node). The Python API also provides functions that return lists of *NodeID*s in a specified order, direction, and set of starting nodes. These traversals are *non-terminating* because they will traverse from the starting nodes until a leaf or root is encountered (depending on the direction).
2. Visitor-based graph traversals are done by creating a visitor that gets "called back" on every node that is visited. These traversals *can be terminated*, by returning a particular value from the visitor callback. This can be advantageous when, for example, the goal is to search for the first encountered node with a particular property.

We provide three graph traversal orders: depth-first search (DFS), breadth-first search (BFS), and topological (Kleinberg and Tardos, 2006). All three can either follow up edges (leaves to roots) or down edges (roots to leaves). The DFS order starts at a particular set of nodes (the "seeds") and visits every node twice, if it is reachable from the seeds. The first visit is in the forward direction and the second visit is after all children of the node have already been visited twice. This DFS traversal is achieved using a last-in-first-out (stack) data structure. The BFS traversal is achieved using a first-in-first-out (queue) data structure. This produces an order where sibling nodes (in tree parlance) are visited closely together. The topological order is equivalent to the DFS order if the entire graph is traversed. Both of these orders impose the topological property: when visiting node $i$ (the second visit, for DFS) it holds that all children of $i$ (nodes reachable from it) have already been visited. The difference is that, for a DFS order, it starts at a set of seeds, traverses the edges, and then visits the nodes (the second time) on the way back "up" to the seeds. The topological order only visits the nodes once, in the "forward" direction from the seeds, but maintains the topological property on *the first visit*. When performing a traversal, one needs to know the seed set to start with: for DFS order the seeds are traversed away from and back to, and only the "back to" maintains the topological property. The topological order traverses away from the seed set and the topological property is maintained for that traversal.

### Constructing a GRG

Given a dataset, let $S = \{0 \ldots N\text{-}1\}$ be the set of all samples and $M = \{M_0 \ldots M_V\}$ be the set of all mutations. Let $S_i$ refer to the subset of samples from $S$ that contain the $i$-th mutation, $M_i$. We'll use $\mathcal{G}$ to refer to a corresponding GRG. The GRG construction problem is to search from the space of all possible internal nodes to find an optimal set of internal nodes such that 1) all mutations in $M$ are correctly mapped to a node and 2) the total number of edges in $\mathcal{G}$ is minimized. Because there are a lot more edges than nodes, the number of edges of $\mathcal{G}$ is an approximation for its size. To make this problem tractable, we split it into two heuristic algorithms: *BuildShape* and *MapMutations*.

*BuildShape* constructs an initial graph $\mathcal{G}$ without any mutations. The goal is to capture haplotype similarity between samples: the more similar two samples' haplotypes are, the more likely those samples will be in the same subgraph of $\mathcal{G}$. In an arbitrary order, we iterate over a set of nodes $S'$ which initially consists only of all the sample nodes (i.e., $S' = S$). This set $S'$ can also be thought of as the roots of a forest of trees, where at each step we merge two trees by creating a new root node and attaching two existing trees as children. To do this, we take each sample $i$ in turn and find the sample $j$ with the most similar haplotype. We create the union of these samples by constructing a new node $N_k$ and making it the parent of nodes $i$ and $j$. This reduces the number of trees in the forest (since we merged two) and the number of nodes in $S'$ (since we remove nodes $i$ and $j$, and add the new node $N_k$) by one. Each node is only considered once, and newly created nodes are also considered in this way. We use Hamming distance to determine haplotype similarity. When comparing internal nodes for similarity with other internal nodes or sample nodes, we use the intersection of their children's mutation sets (in practice, we treat the minor alleles in a dataset as mutations). In other words, internal node similarity is measured against the consensus haplotype of the descendants of that node.

The output of a single *BuildShape* invocation is a binary tree in GRG format called a tGRG. A single tree obviously could not capture the relationships between all mutations and samples, so we have a parameter $T$: the genome is split into $T$ segments and *BuildShape* constructs a GRG tree for each. Experimentally, it is optimal to split the genome into segments of size 50-150Kbp for most non-simulated datasets. The $T$ GRG binary trees that cover the genome are then merged into $P <= T$ sub-GRGs. For simplicity, the description in the main text assumes $P = T$. $P$ determines how parallelizable GRG construction is: each merged sub-GRG can have mutations in the corresponding genomic region mapped onto it in parallel. In practice, $P$ is expected to be larger than the number of cores available for GRG construction.

### Parameter P

$P$ determines how parallelizable GRG construction is: each merged sub-GRG can have mutations in the corresponding genomic region mapped onto it in parallel. An increase in $P$ only marginally increases the size of the final GRG but significantly reduces overall runtime even when construction is run on a single core (see **Fig. S3**). A large $P$ also reduces RAM usage per thread of execution, as fewer variants are mapped at a time. The trade-off is that, as $P$ grows, it is more likely that mutations with long-range linkage will not have the opportunity to share common sample subsets (because they are being mapped independently from each other).

### Finding Similar Haplotypes

**Fig. S14** illustrates three different haplotype representations. In the case of a sequence, we assume 1 byte per character. The sparse mutation list only contains differences from the reference genotype, and we assume that the variants (alternative alleles) in the dataset are stored and numbered from 0…($V$-1) so that a mutation list is just a list of integer indexes.

Taking chromosome 17 from the 1000 Genomes Project (The 1000 Genomes Project Consortium et al., 2015) as an example: $V$ = 2,341,841 and on average each sample deviates from the genome at ~82,000 sites (approximately 3.5%). This means the sequence storage would be 2,341,841 bytes (2.23MB) per sample, the sparse list would be $(0.035 \times 2341841 \times 4) \sim= 327{,}858$ bytes (320KB) per sample.

Consider a case where we want to scale up to a million haplotypes (500,000 diploid individuals): with such a dataset, the sparse representation would still use more than 300GB of memory. Furthermore, the sparse representation is not fixed-length per sample, which complicates and slows down Hamming distance calculations. To further overcome this limitation, during GRG construction, we use a simple bloom filter representation (**Fig. S14c**), with $B = 4$ bits per mutation (on average). This allows us to construct a high-quality GRG (**Fig. S15**) but uses 8 times less memory than mutation lists, getting us down to 40982 bytes (40KB) per sample. Our million haplotypes can now fit in approximately 38GB of RAM.

The two computations that we need to do on our haplotypes can also be done very efficiently on the bloom filter representation:

1. When merging root nodes during tGRG construction (*BuildShape*) we need to intersect the haplotype children of the node. This is just the bitwise AND operation across all $m$ bits of the bloom filter.
2. When searching for nearest neighbors we need to compute the Hamming distance between two haplotypes. This can be done by XORing the bloom filter representation and then counting the bits that are set (1s) after XOR. The *popcount* instruction can be used on x86/64 hardware, but we've found that the software implementation in **Algorithm S1** is slightly faster.

```
uint32_t xorValue = filter1[i] ^ filter2[i];
static const int S[] = {1, 2, 4, 8, 16}; // Magic Binary Numbers
static const int B[] = {0x55555555, 0x33333333, 0x0F0F0F0F, 0x00FF00FF,
                        0x0000FFFF};
uint32_t hammingDist = xorValue - ((xorValue >> 1) & B[0]);
hammingDist = ((hammingDist >> S[1]) & B[1]) + (hammingDist & B[1]);
hammingDist = ((hammingDist >> S[2]) + hammingDist) & B[2];
hammingDist = ((hammingDist >> S[3]) + hammingDist) & B[3];
hammingDist = ((hammingDist >> S[4]) + hammingDist) & B[4];
```

**Algorithm S1**: C implementation of Hamming distance between two 32-bit values using "Counting bits set, in parallel" from the "Bit Twiddling Hacks" site (https://graphics.stanford.edu/~seander/bithacks.html)

We also build a nearest-neighbor index using a BK-tree data structure (Burkhard and Keller, 1973) to speed up the similarity search. A BK-tree enables sublinear-time search (on average), provided that the distance measure used is 1) discrete and 2) metric (satisfying the triangle inequality, among other properties). Nodes in the BK-tree are data points (haplotypes in our case) that can have one or more items (samples) associated with them (**Fig. S16**). Child edges are labeled with the metric distance to the data point associated with the child node. For any node $n$, every descendant reachable from the edge labeled $D$ is exactly distance $D$ away from the datapoint associated with $n$. We search for the nearest neighbor of our query sample $S_2$ by computing the distance between $S_2$ and each node of the tree. To traverse from a parent node to a child node, we can use the edge labels and reverse triangle inequality to rule out some subtrees, as they are guaranteed to not have a nearer neighbor than the current nearest (**Fig. S16**).

### Mutation Mapping

We map mutations onto the graph losslessly: every mutation node $M_i$ should transitively reach the set of sample nodes $S_i$ and no sample nodes apart from $S_i$. If the existing graph shape matches our mutations nicely then we will reuse a lot of existing nodes. There are three possible scenarios (depicted in **Fig. S2**), the two more common ones require creating a new node to represent the mutation. In the worst case, there are no reusable nodes in the graph and we need to create an edge from the node containing $M_i$ to each sample node in $S_i$ (**Fig. S2b**), which is equivalent to storing the mutation as a sparse mutation list.

Mutations are mapped using **Algorithm S2**. Intuitively, for a mutation $M_i$ we do an upwards search from the sample nodes representing $S_i$ and keep a counter of how many samples from $S_i$ reach each internal node. If a node has more samples in general than this counter, it means there are samples beneath it that are not in $S_i$ (and thus it is unmappable). We terminate the graph search when this occurs, and save all of the nodes that we traversed immediately before the terminating node. These candidate nodes are potentially reusable for mapping $M_i$ and after searching the graph we greedily use each candidate in descending order of covered sample set size. Candidates with overlapping covered samples are not allowed.

In **Algorithm S2**, *depthFirstOrder* is the list of all nodes in the GRG, sorted by a post-order, depth-first search. *topologicalOrder* is sorted the same way but only lists the nodes that are reachable via up-edges from the given list of starting nodes. Sets of pairs are treated like maps from the first item in the pair to the second. The [ ] (square bracket) notation is used to lookup/modify an item in sets of pairs. *sortByLargestCoveringSet* takes a set of nodes and returns them as a list where the first node covers the most samples and the last node covers the least samples. *getCoveredSamples* retrieves the covered samples, as a set, for a given GRG node.

**Algorithm S2** Map mutations to an existing GRG

```
mapMutations(grg, mutationList):
  sampleCountMap ← ∅
  for all node ∈ depthFirstOrder(grg) do
    samplesBeneath ← 0
    for all childNode ∈ node.downEdges do
      samplesBeneath ← samplesBeneath + sampleCountMap[childNode]
    end for
    sampleCountMap[node] ← samplesBeneath
  end for
  for all mutation ∈ mutationList do
    candidateNodes ← ∅
    mutationSampleMap ← ∅
    for all node ∈ topologicalOrder(grg, mutation.sampleNodes) do
      if node ∈ mutationSampleMap ∨ node.isSample then
        samplesBeneath ← 0
        possibleCandidates ← ∅
        for all childNode ∈ node.downEdges do
          if childNode ∈ mutationSampleMap then
            samplesBeneath ← samplesBeneath + mutationSampleMap[childNode]
            possibleCandidates ← possibleCandidates ∪ childNode
          end if
        end for
        if samplesBeneath ≠ sampleCountMap[node] then
          candidateNodes ← candidateNodes ∪ possibleCandidates
        else
          mutationSampleMap[node] ← samplesBeneath
        end if
      end if
    end for
    mutationNode ← makeNode(grg)
    mutationNode.mutations ← {mutation}
    alreadyCovered ← ∅
    for all candidateNode ∈ sortByLargestCoveringSet(candidateNodes) do
      sampleSet ← getCoveredSamples(candidateNode)
      if sampleSet ∩ alreadyCovered = ∅ then
        alreadyCovered ← alreadyCovered ∪ sampleSet
        addEdge(grg, mutationNode, candidateNode)
      end if
    end for
    for all uncoveredSampleNode ∈ (mutation.sampleNodes − alreadyCovered) do
      addEdge(grg, mutationNode, uncoveredSampleNode)
    end for
  end for
```

### Merging GRGs

The GRG merging algorithm (**Algorithm S3**) proceeds by modifying *grg1* until it contains the same mutation-to-sample relationships as both *grg1* and *grg2*. We consider nodes equivalent if they cover the same samples. Nodes in *grg1* are reused for *grg2*'s mutations if they are equivalent. Otherwise, the *grg2* nodes (and its edges) are recursively copied to *grg1*. The *oneWayHash* function captures this equivalence by hashing a set of GRG nodes with an algorithm that is extremely unlikely to have unintended collisions. In our implementation, we use the MD5 hash (Rivest, 1992).

**Algorithm S3** Merge two GRGs

```
grgMerge(grg1, grg2):
  hashToNode ← ∅
  for all node ∈ depthFirstOrder(grg1) do
    hashToNode ← hashToNode ∪ (node, oneWayHash(getCoveredSamples(node)))
  end for
  grg2ToGrg1Map ← ∅
  for all node ∈ depthFirstOrder(grg2) do
    hashValue ← oneWayHash(getCoveredSamples(node))
    if hashValue ∈ hashToNode then
      nodeInGrg1 ← hashToNode[hashValue]
      nodeInGrg1.mutations ← nodeInGrg1.mutations ∪ node.mutations
    else
      nodeInGrg1 ← makeNode(grg1)
      nodeInGrg1.mutations ← node.mutations
      for all child ∈ node.children do
        addEdge(grg1, nodeInGrg1, grg2ToGrg1Map[child])
      end for
    end if
    grg2ToGrg1Map ← grg2ToGrg1Map ∪ {(node, nodeInGrg1)}
  end for
```

After merging GRGs into a single graph, we simplify the graph and then serialize using the Compressed Sparse Row (CSR) format (Duff, 1984). **Fig. S12** illustrates this format for a simple graph. CSR format stores all edges in a single linear array, and a separate array (the edge index array) maps a given node *i* to its starting position in the edge array. The edges for node *i* are between $edgeIndexArray_i$ (inclusive) and $edgeIndexArray_{i+1}$ (exclusive). If $edgeIndexArray_i == edgeIndexArray_{i+1}$, then node *i* has no edges. For convenience, there are *n*+1 entries in the edge index array for *n* nodes: the last entry will always be the size of the edge array (the total number of edges in the graph).

Since the GRG is fully described by its down edges (its up edges are just inverted down edges) we only serialize the down edges. Deserialization is then slightly more expensive, as we have to reconstruct the up edges while reading the CSR data. When a GRG is immutable it is stored in memory in CSR format (Duff, 1984), since it is extremely compact and much more cache-efficient than the alternative (edge lists). All GRGs are stored in CSR format when on disk.

### Indexable Genotype Data (IGD)

Parallel construction of GRG requires being able to quickly load haplotype data from a file starting at a particular genomic position. Text-based formats like VCF are inefficient with this somewhat random file access, so we create a simple, compact, *indexable* format called “IGD” that we use during GRG construction. GRG can still be created directly from VCF (or compressed VCF), but it is quite slow. It is significantly faster to first convert the VCF to IGD and then construct a GRG (**Fig. S6b**). BGEN is also quite fast when used as input to GRG construction, but the conversion of VCF to BGEN for *phased data* is quite slow (**Fig. S6a**).

We have released an open-source, lightweight VCF parser and IGD library that allows easy conversion between VCF and IGD, and supports read IGD files (https://github.com/aprilweilab/picovcf). The file format itself is binary (not text-based), and expands multi-allelic variants to be binary: a single variant with *k* alternate alleles becomes instead *k* variants each with a single alternate allele. This closely matches the definition of "mutation" that we use through GRG. A more detailed description of the file format can be found at https://github.com/aprilweilab/picovcf/blob/main/IGD.FORMAT.md.

### File Format Conversions

**Table S1** gives a breakdown of various popular file formats for genome sequences compared against the GRG and IGD file formats. Notably, the GRG compressed the data significantly more for the large sample size data (UK Biobank) than the very small sample size data (1000 Genomes Project).

Currently, GRG does not support unphased data, but there is nothing fundamental about the graph encoding that prevents storing unphased data. We expect to support unphased data in future work.

**Fig. S6c** gives a breakdown of various file format conversion operations and their file sizes on a subset of the 1000 Genomes Project dataset. Building a GRG directly from *vcf.gz* files is quite slow, both because of the overhead of decompressing and parsing the text data, and because we are not using indexed *vcf.gz* files (e.g. via Tabix files (Bonfield et al., 2021) to skip to the part of the genome for which we are constructing tGRGs.

We often compare with both *vcf.gz* and VCF for file *sizes*, because the uncompressed size of the VCF file gives an idea of how much text data needs to be parsed when using a VCF file (whether it is compressed or not). However, for large datasets the *vcf.gz* always needs to be used directly, because the uncompressed VCF file would not fit on most hard drives.

Some of the meta-data that is often stored in VCF files includes sample identifiers, variant identifiers, and quality metrics for a particular variant. We believe this metadata is best stored separately from the GRG (and IGD) files for two reasons. First, they are all orders of magnitude smaller than the actual genotype data, and thus need no special storage or processing. Second, it may be desirable to examine or edit this data manually, so storing them in a standard text format (such as tab or comma-delimited files) is much more convenient than a binary format.

### Comparison of file sizes

We compared GRG against XSI and Savvy for file size, time to compress, and a handful of computations. Both XSI and Savvy are based on BCF, which means that they support meta-data and non-genotype data that GRG does not. Thus for a fair comparison we stripped all BCF files of non-GT data.

Both XSI and Savvy use two techniques for compression: sparse representation for low-frequency variants, and PBWT sorting with zstd compression for high-frequency variants. Additionally, XSI uses word-aligned hybrid (WAH) to encode the data prior to zstd compression.

We used xSqueezeIt (XSI) v4.0.1 and Savvy (SAV) v.2.1.0.

*Stripping BCF file of non-GT contents:* `bcftools annotate --remove FORMAT <input bcf> -O b --output <output bcf>`

*Generating an XSI file using xSqueezeIt:* `xsqueezeit -c -f <input vcf> -o <output xsi>`

*Generating a SAV file using Savvy:* `sav import --pbwt-fields GT --sparse-threshold 0.01 --phasing full <input bcf> <output sav>`

To project the size of uncompressed VCF files, we constructed simulated datasets of different numbers of variants and samples. We then used linear regression to find a function of the number of variants ($x$) that predicts the megabytes per sample. This gave us the equation: $(1.39 \times 10^{-6})x + 0.234$. For example, solving for UK Biobank chromosome 2 (which has approximately 7,720,000 variants) gives us 10.97 MB / sample. Since we have 400,011 samples in that dataset, we get a total projected VCF size of 4.18TB.

### Benchmark for Allele Frequencies

To evaluate the computational efficiency of calculating genome-wide allele frequencies across different data structures and algorithms, we used tabular data structures (VCF and bed files) within plink v.1.90b6.26 and our GRG structures as input. The Phase III data in the 1000 Genomes Project is used for benchmarking https://hgdownload.cse.ucsc.edu/gbdb/hg19/1000Genomes/phase3/. Allele frequencies of 22 autosomes from the 1000 Genomes Project were computed using the *plink.freq* function and the *grgl.freq* function respectively (**Fig. S5c**). All computations were executed using a single computational CPU thread for all chromosomes and across both data structures.

We also compared allele frequency calculations between GRG, tskit, XSI, and Savvy, using the same simulated dataset we used for measuring GRG construction times/sizes. For XSI, we used a modified version of the *af_stats* example they provide in their repository. The modification was to output a text file instead of a BCF file, in order to provide a more fair comparison to the other tools (we expect that outputting BCF would be *more* expensive than a text file). We also created a separate executable, based on *af_stats*, that operates on a subset of the genome (a region) to compare against Savvy and GRG. For Savvy we wrote utilities to calculate allele frequency on the entire file, a subset of the genome (region), and a subset of the samples. We used *savvy::reader* to read each *savvy::variant* and then *savvy::variant::get_format()* to get the genotype data for each variant and compute the allele frequency. We used *savvy::reader::subset_samples()* to restrict the samples for allele frequency computation, and *savvy::reader::reset_bounds()* to restrict to a subset of the genome. For tskit we used *TreeSequence.sample_count_stat()* to calculate allele frequencies for the entire genome.

As far as we could tell from the XSI documentation it does not support fast indexing by a subset of samples. In **Fig. S10d** we can see that GRG is significantly (5-90x) faster than Savvy when computing allele frequency for a subset of samples.

It is worth noting that Savvy and XSI are both tabular formats with indexes: XSI supports fast indexing based on genome position (regions), and Savvy supports fast indexing by both position and sample IDs. To explore their behavior in a subset of samples or a subset of regions, we tested allele frequency computations in three sub-regions, with length of 0.1Mbp, 1Mbp and 5Mbp with variable sample sizes. When the size of the sub-region of interest increases to 5Mbp (**Fig. S10**), GRG is faster than both XSI and Savvy, with the performance gap widening in larger sample sets. We expect this is because XSI and Savvy make use of explicit compression (zstd), which requires decompression before computations can proceed. The decompression time can significantly affect the total computation time, especially when sample size or sequence length increase. In contrast, the compression in GRG is implicit in its data structure: there is no decompression step needed and computations can be implemented directly on the original data form.

### Zygosity Information Computation

Another example of a GRG graph computation is the zygosity information for each variant. The goal is to calculate, for each variant, the number of diploid individuals with (a) zero copies of the alternate allele, (b) one copy of the alternate allele, and (c) two copies of the alternate allele. Similar to the allele frequency and GWAS calculations, we traverse the graph once in topological order and propagate the values of interest from child node to parent node. At each node we store the pair of values *(samples_beneath, homozygous_beneath)* which represent the number of haploid samples beneath the node, and the number of individuals that have already coalesced at or beneath the node. Every sample node is initialized with the pair *(1, 0)*. We traverse every node in topological order and sum the values of its children to obtain the current node's values. Additionally, the GRG itself stores *numIndividualCoals* at each node, which is the number of individuals that coalesce *exactly* at that node. Each node's update pseudo-code is then:

```
samples_beneath[node] = 0
homozygous_beneath[node] = node.numIndividualCoals
for child in get_children(node):
  samples_beneath[node] += samples_beneath[child]
  homozygous_beneath[node] += homozygous_beneath[child]
```

After traversing the whole graph we can compute the results for variant at node *variant_node* as:

```
heterozygous = samples_beneath[variant_node]/2 - homozygous_beneath[node]
homozygous = homozygous_beneath[node]
nocopies = totalIndividuals - (heterozygous + homozygous)
```

### GRG as a Reduction from ARG

The term ARG has been used to describe several different things in the literature (Wong et al. 2024). Here we describe the *information* that is removed from an ARG to produce a GRG, but our description makes the most sense when considering the genome ARG (gARG) *representation* of an ARG. Related to our study, *true* ARGs are equivalent to the underlying real genealogical history of a sample set. In simulations (Baumdicker et al., 2022), the paths of inheritance and coalescence times are often simulated first, and then neutral mutations are overlaid on top of these paths to generate haplotypes. Here we also refer to the actual paths of inheritance and coalescence as *true* ARG, and the generated haplotype data as *true* data. Mutation events, by definition, can always be overlaid to a true ARG to encode the true data, but not all parts of a true ARG bear mutation events. The GRG could be regarded as a more compact ARG-like graph that only captures the paths used for the actual "inheritance" of mutations to a set of haplotypes. The *optimal* GRG could be defined as a reduction from the true ARG in four aspects (**Fig. 5b**), because it is the GRG that captures the haplotype formation history. First, in order to record the coalescence time, the same topological structure differing only by coalescence times must be represented distinctly in an ARG. These duplicated structures can be omitted in a GRG because GRG does not encode coalescence times. Second, paths in a true ARG that do not serve the purpose of passing down any mutations can be removed. Third, when two or more mutations are inherited by the same set of haplotypes through different paths, these paths could be merged. Fourth, edges that are never used as the first edge in any mutation path could be removed. After doing this reduction, we get a new graph that compactly encodes the true genotypes. If we encode this new graph using the GRG data structure, we get an *optimal* GRG that captures the haplotype evolution history.

### Tree Sequence to GRG Conversion

Tree sequences (an ARG format) are converted to GRGs by copying and duplicating nodes and edges. In **Algorithm S4**, *markMRCA* finds the most recent common ancestor (MRCA) of a set of nodes and returns the set of nodes on the path to the MRCA. *makeNode* constructs a new GRG node. *mutation.node* is the tree sequence node associated with (beneath) a given mutation. *treeNodeToGrgNode* is a set of pairs that maps tree sequence nodes to GRG nodes.

**Algorithm S4** Construct a GRG from Tree-Sequence

grgFromTreeSeq(treeSequence):
  $grg \leftarrow \emptyset$
  $splitNodes \leftarrow \emptyset$
  $treeNodeToGrgNode \leftarrow \emptyset$
  **for all** $tree \in treeSequence$ **do**
    $impactedNodes \leftarrow \emptyset$
    **for all** $edgeParent, edgeChild \in tree.edgeDiffsFromPrevious$ **do**
      $impactedNodes \leftarrow impactedNodes \cup (edgeParent, edgeChild)$
    **end for**
    $splitNodes \leftarrow splitNodes \cup markMRCA(impactedNodes)$
    **for all** $mutation \in tree.mutations$ **do**
      $grgNode \leftarrow addMutationFromTree(grg, mutation.node, splitNodes, treeNodeToGrgNode)$
      $grgNode.mutations \leftarrow \{mutation\}$
      $grg \leftarrow grg \cup \{grgNode\}$
    **end for**
  **end for**
  **return** $grg$

addMutationFromTree(grg, treeNode, splitNodes, treeNodeToGrgNode):
  $canReuse \leftarrow treeNode \in treeNodeToGrgNode$
  $canReuse \leftarrow canReuse \wedge (treeNode.isSample \vee (treeNode \notin splitNodes))$
  **if** $canReuse$ **then**
    **return** $treeNodeToGrgNode[treeNode]$
  **end if**
  $splitNodes \leftarrow splitNodes - treeNode$
  $newParentNode \leftarrow makeNode(grg)$
  **for all** $child \in treeNode.children$ **do**
    $newChildNode \leftarrow addMutationFromTree(child, splitNodes, treeNodeToGrgNode)$
    $newParentNode.children \leftarrow newParentNode.children \cup \{newChildNode\}$
    $treeNodeToGrgNode \leftarrow treeNodeToGrgNode \cup (treeNode, newParentNode)$
  **end for**
  **return** $newParentNode$

### The Robustness of GRG Construction

To test the optimal size of GRG, we performed *msprime* simulations (Baumdicker et al. 2022) to generate *true* ARGs and *true* data (see **Methods**). ARGs can be compactly encoded with the tree sequence format (Kelleher et al. 2018). We construct GRGs in two different ways. First, we construct GRGs from true ARGs to investigate the size of the *optimal* GRG. To this end, we find that the optimal GRG is consistently the most compact data structure across sample sizes. Moreover, the size of optimal GRG grows sublinearly with sample size (**Fig. S7ab**).

Next, we investigate whether we can obtain a more compact GRG by first inferring an ARG. In this experiment, true data are generated by *msprime* simulation (see **Methods**). While ARG-Needle (Zhang et al. 2023) is one of the more scalable ARG inference techniques, the mutation events are not inferred, thus we cannot use it directly for GRG construction. To date, *tsinfer* (Kelleher et al., 2019) is the only ARG inference method that is scalable to at least 10,000 whole genomes that still maps mutation events.

Applying *tsinfer* to true data, we find that the *optimal* GRG (*true* TS-GRG) is smaller than the TS-GRG, which is smaller than the constructed GRG (**Fig. S7**).

To investigate whether the *inferred* TS-GRG is smaller than the constructed GRG in *real* data, we applied *tsinfer* to the 1000 Genomes Project data (The 1000 Genomes Project Consortium et al., 2015). Interestingly, we find that the *inferred* TS-GRG is about two times larger than the constructed GRG (**Fig. S7c**). When genetic data are not accurately ascertained, the true ARG may no longer accurately encode the observed sequences (that contain errors), because errors in the data are not modeled by the true ARG. Therefore, we hypothesize that real data, such as those collected by the UK Biobank and the 1000 Genomes Project, are subject to sequencing and variant calling errors, and ARG inference methods might not perform robustly in the presence of such errors, whereas the similarity search algorithm used in *BuildShape* tolerate such errors better. To this end, we performed a simple test to compare the sizes of the *inferred* TS, *inferred* TS-GRG, and constructed GRG using simulated data that has 0.1% added noise, which roughly mimics the variant calling error in the UK Biobank (Saunders et al.,2007; Browning and Yu, 2009; Halldorsson et al., 2022). Although this noise simulation procedure is oversimplified, we find that the sizes of the *inferred* TS, *inferred* TS-GRG, and constructed GRG all increase significantly as the noise level increases, suggesting that noise can indeed have a sizable influence on graph sizes. Interestingly, the size of constructed GRG becomes consistently smaller than *inferred* TS and *inferred* TS-GRG across a range of sample sizes even when there is only 0.1% of added noise (**Fig. S8ab**), suggesting that our GRG construction algorithm is more robust to errors in the data ascertainment.

We then used the ratio of the number of mutations in the *inferred* tree sequence (TS) to the number of input alternative alleles in the VCF (recurrent mutations are rare, so this is still a good approximation to the number of mutations) as an index to illustrate the actual number of mutation events identified after ARG inference. This index also helps illustrate the levels of ARG inference bias under different simulation settings. Consistent with the file size comparisons, when errors are added to the simulated data, *tsinfer* tends to infer more mutation events (**Fig. S8ef**), resulting in a higher ratio. This robustness to noise in the constructed GRG is likely because our GRG construction decouples the BuildShape and MapMutations steps, hence not making any explicit assumptions about mutation models.

We explore the effects of noise in the dataset by randomly flipping alleles in VCF files produced from *msprime* simulations. For a given percentage of noise, $P_n$, and a dataset with $N$ individuals and $V$ variants, we uniformly sample $(P_n / 100) \times N \times V$ pairs of (individuals, sites) and flip each allele from its current value in the set {0|1, 1|0, 1|1, 0|0} to one of the other values in that set. We chose our experimental value of $P_n$ = 0.1% to be in line with common error estimates on medium to high coverage DNA sequencing (i.e., 20-30×) data.

### Simulations with a Realistic Recombination Map

Numerous studies have shown that the recombination rates vary along the genome and play a significant role in shaping the local density of genetic variants within the population (McVean et al., 2004; Pratto et al., 2014). Thus, to better represent the empirical data in simulations, we incorporate a realistic recombination map from the 1000 Genomes Project (The 1000 Genomes Project Consortium et al., 2015), using the first 100Mbp of chromosome 1 to the simulation framework, and simulated 100,000 diploid individuals. We find that the resulting number of genetic variants and size of constructed GRG from this simulation are almost the same as those of a simulation with a flat recombination rate (**Fig. S17**).

### Simulations with Human Demographic History

Most of the empirical sequencing data in the UK Biobank are non-Africans, with a demographic history involving population bottleneck and migration. To better resemble the empirical data, we also extend the simulation framework with a finer-scale demographic history model and gene conversion (**Fig. S17**) using stdpopsim v.0.2.0 (Adrion et al., 2020). The demographic model is adapted from (Jouganous et al., 2017) to represent a four population out-of-Africa history and we only sample central European haplotypes in the output. The full human chromosome1 is simulated with the realistic recombination map from the Phase II HapMap project (The International HapMap Consortium, 2007). Gene conversion fraction is set to 0.02 with a length of 300bp based on the parameter choices of a recent study (Browning and Browning, 2023). With this simulation framework, the number of mutations is about 7-fold higher compared to the simple simulation with a flat recombination rate, and the resulting size of the constructed GRG (947MB with $T = 500$) is roughly 4.67-fold larger (compared to 203MB with $T = 100$). Interestingly, when the density of genetic variants is low, increasing the parameter $T$ does not lead to further compression of the constructed GRG. In contrast, the opposite effect is observed in the simulations with high variant density, consistent with our results in **Fig. S3**. Based on this result, we adopt a higher $T$ when constructing GRGs from the 1000 Genomes Project and the UK Biobank data.

### Graph Statistics

**Fig. S18** tries to help answer two questions: “how similar is the GRG hierarchy to the corresponding ARG?” and “how optimal is the compression provided by the current GRG construction?”. These two questions are highly related, since reducing an ARG to a GRG should provide the near-optimal compression. We compared the true TS-GRG and the constructed GRG for simulated data for 100,000 individuals. Here the true TS-GRG means we simulated data using *msprime*, which emitted a ground-truth ARG in tree sequence format, and then reduced that ARG to a GRG via the tree sequence to GRG algorithm (see **Algorithm S4**). As **Fig. S18** illustrates, this “true TS-GRG” has significantly more hierarchy than the constructed GRG: the former has many more nodes, and fewer edges per node. **Fig. S18ab** indicate that the true TS-GRG has fewer edges per node, many more nodes, and each node on average has many internal nodes beneath it. Similarly, panels **c** and **d** demonstrate that nodes in the true TS-GRG have lower *local utility*, but higher *global utility*. For global utility, it is a bit more nuanced: the true TS-GRG has a more even distribution of utilities, whereas the constructed GRG has a lot of nodes that do the majority of the “heavy lifting.” I.e., the peak on the right hand side of panel **d** represents the nodes that capture the majority of reuse/compression for the graph. These statistics seem to indicate that there is room to improve the GRG construction algorithm to capture more hierarchy, and thus more compression.

### Supplementary Tables

**Table S1: Comparison of different file formats for aligned whole-genome sequencing data.**

| Format | Needs decompression | Text-based | Phased/Unphased | Avg size multiplier over GRG (1,000 Genomes Project) | Avg size multiplier over GRG (UK Biobank) |
|---|---|---|---|---|---|
| VCF | No | Yes | Both | 135.9× | 2000× *(estimated)* |
| *vcf.gz* | Yes | Yes | Both | 2.3× | 12.7× |
| BGEN | Yes (typically) | No | Both | 2.7× | N/A* |
| PLINK BED, BIM, FAM | No | Partially | Unphased | 8.4× | 179.9× |
| IGD | No | No | Both | 2.3× | 3.7× |
| GRG | No | No | Phased | 1.0× | 1.0× |
| XSI | Partial | No | Both | Not calculated | 0.47x † |
| Savvy | Yes | No | Both | Not calculated | 0.56 † |

*Not estimated because conversion to phased BGEN files does not have scalable tool support. The time and cost of conversion would have been prohibitive.

† Only chromosomes 13 and 22 were compressed with XSI and Savvy, due to the cost of first converting to BCF format.

**Table S2: Comparison of different file formats for the SARS-CoV-2 sequences.**

| Date | Samples | Size (.ts.tsz) | Size (.ts) | Size without metadata (.ts) | Size (.grg) | Construction time |
|---|---|---|---|---|---|---|
| Up to June 30, 2021 | 1,270,000 | 58MB | 891MB | 47MB | 37MB | 5.4 seconds |
| Up to June 30, 2022 | 657,000 | 37MB | 480MB | 38MB | 31MB | 3.6 seconds |

### Supplementary Figures:

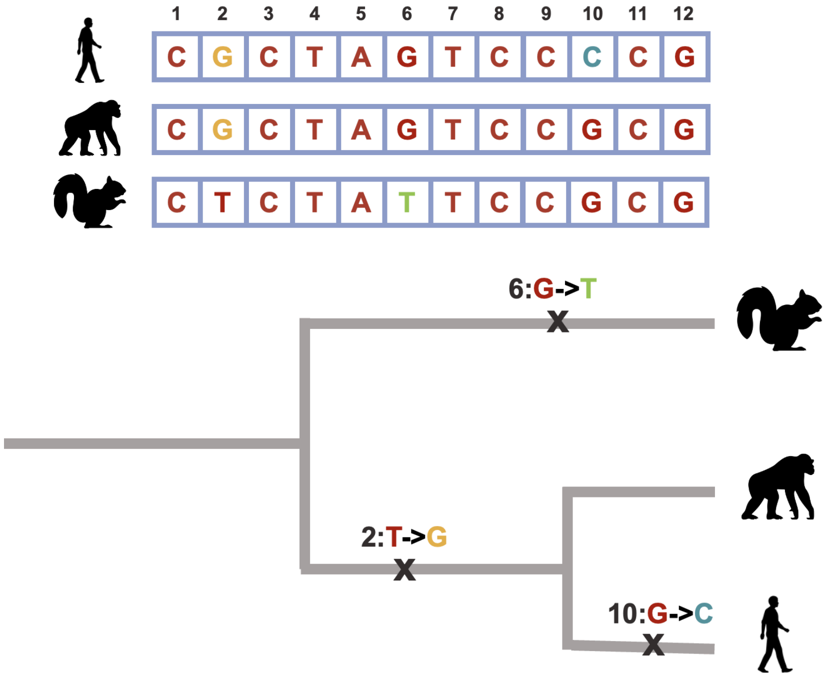


**Figure S1: Using a phylogenetic tree to encode DNA sequence alignment.** Here, we present a hypothetical example of using a phylogenetic tree for encoding the aligned genomes across three species: human, chimpanzee and squirrel. In the upper panel, a section of the aligned genomes of these species is shown, with ancestral alleles highlighted in red and the derived alleles across species shown in other colors. To represent the sequence alignment using a phylogenetic tree, only the mutation events that generate the derived alleles need to be specified in the phylogenetic tree, as all ancestral alleles are shared across three species and are fixed. The mutation events contributing to the derived alleles are labeled along the branches of the phylogenetic tree.

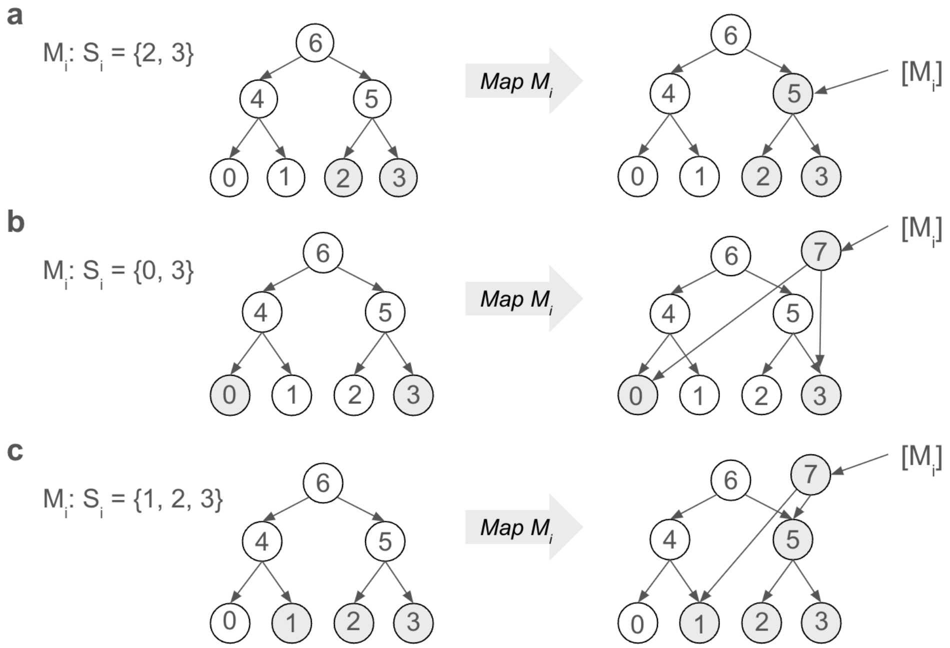


**Figure S2: Three example scenarios when mapping a mutation $M_i$ to a GRG. a:** All the samples for $M_i$ are beneath a single reusable node 5, so we just associate $M_i$ with that node. **b:** There are no reusable nodes (aside from sample nodes) for $M_i$, create new node 7 and singleton edges to the needed samples. Associate $M_i$ with new node 7. **c**: There is a reusable node 5, but it only captures 2/3 of the needed samples. Create a new node 7, an edge to sample 1, and an edge to reusable node 5. Associate $M_i$ with new node 7.

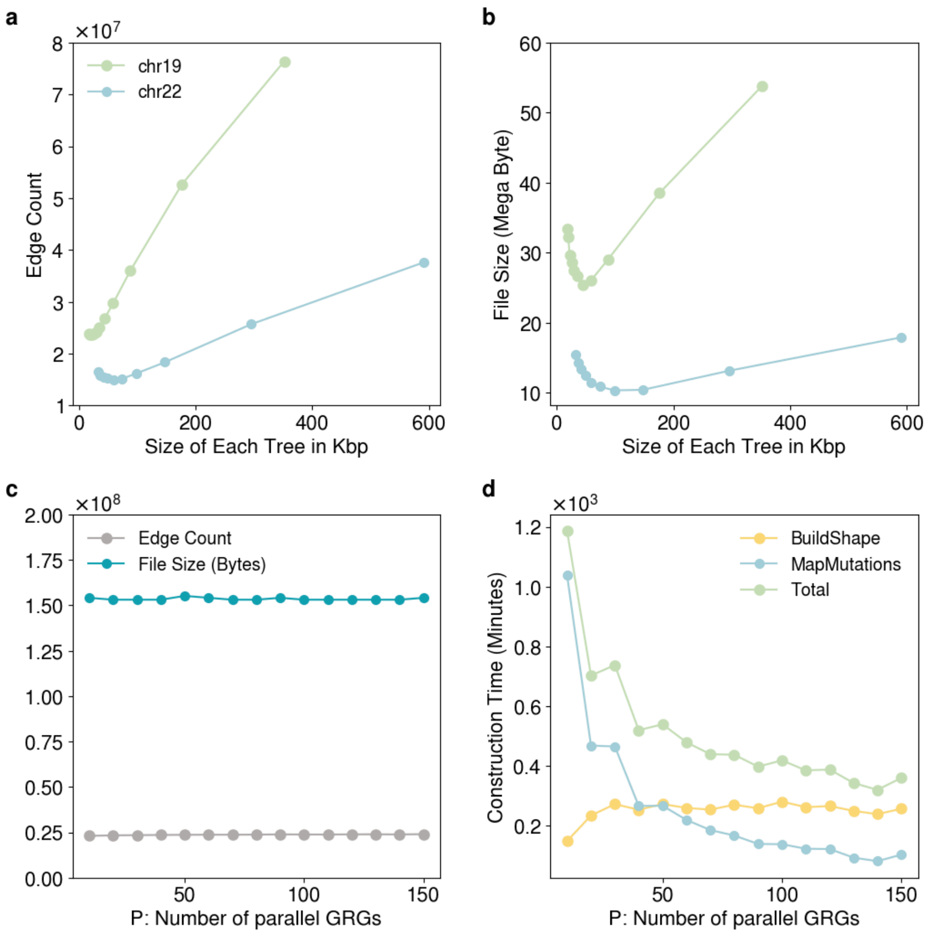


**Figure S3: Parameters *T* and *P* control the performance of GRG construction. a-b**: Parameter *T* controls how many tree-GRGs to construct before mapping mutations. We plot the segment length used for each tree, which can then be converted into the parameter *T* using the chromosome length. Having too few trees has a significant impact on graph size. After a certain point, having too many trees slows down GRG construction and no longer reduces graph size. Data is from chromosomes 19 and 22 of the 1000 Genomes Project dataset. **c-d**: The majority of mutation correlations maintain locality, so increasing *P* has a very minor impact on the final graph size. Up to a point, increasing *P* reduces CPU time significantly. It further reduces *elapsed* time when there are multiple cores for parallelization (not shown in figures). Data is from chromosome 19 of the 1000 Genomes Project dataset.

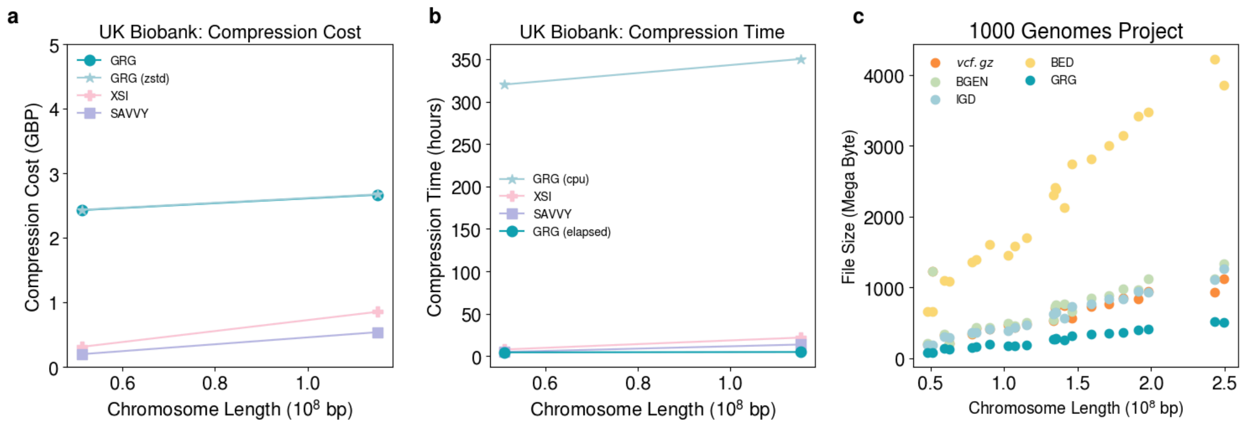


**Figure S4: Comparison of file formats for aligned whole-genome sequencing data on the real datasets. a**: compression time for converting the whole-genome polymorphisms of chromosome 13 (right) and chromosome 22 (left) from tabular formats (i.e. BGEN or BCF) to GRG, XSI and Savvy on the UK Biobank dataset. **b**: Compression time for different formats conversion. See **Table S1** for a summary of file format features. **c**: Comparisons of file sizes for encoding whole-genome polymorphisms in the 1000 Genomes Project dataset.

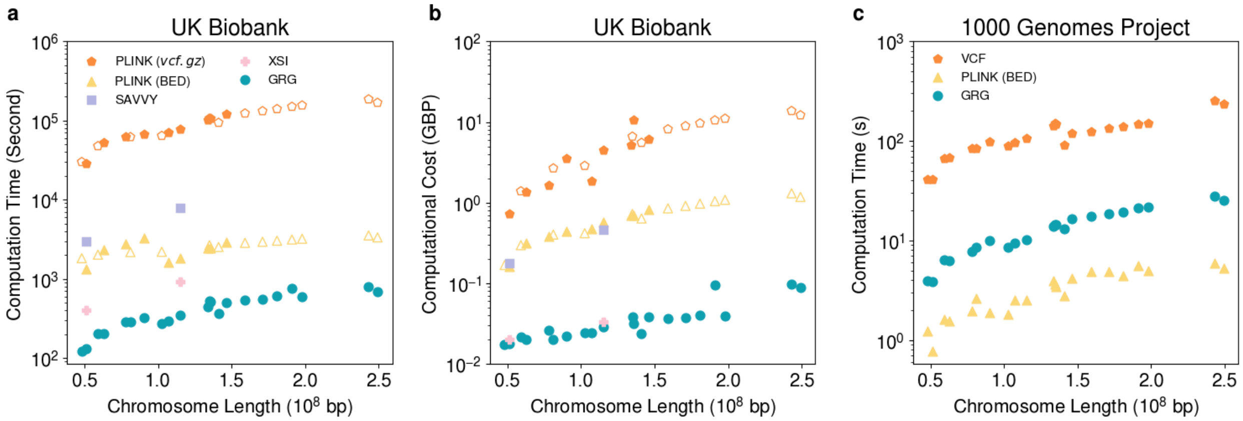


**Figure S5: Allele frequency computation.** **a**: comparison of computational efficiency in the UK Biobank. Allele frequencies were computed on 9 chromosomes (chromosomes 8, 10, 12, 14, 16, 18, 20, 21, 22, labeled as solid dots). For Savvy and XSI, we focused on 2 chromosomes (chromosome 20 and 22). These results were used to predict the expected computation time for the remaining chromosomes (presented as open dots) with a linear projection. For Savvy and XSI, we focused on 2 chromosomes (chromosome 13 and 22). **b**: Comparisons of computational costs in Pound Sterling (GBP) for calculating genome-wide allele frequencies in the UK Biobank Research Analysis Platform. Similar to **a**, solid dots represent the results from our experiments while open dots represent the projected computational cost for the remaining chromosomes. **c**: comparison of computational time for allele frequencies in the 1000 Genomes Project: each data point represents the computational time of genome-wide allele frequencies for each chromosome in the 1000 Genomes Project, either using tabular data structures (VCF and plink bed files (PLINK)) or GRG structure as input.

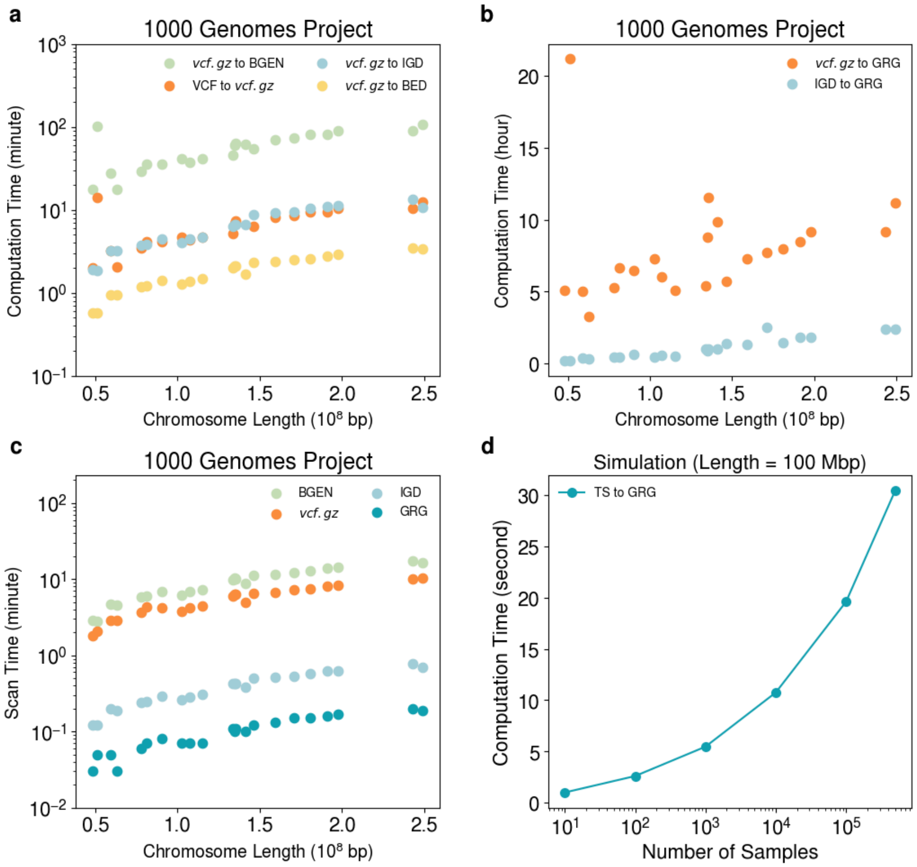


**Figure S6: Comparison of file formats for phased whole-genome sequencing data. a-c** are the results for the 1000 Genomes Project where each dot represents a full chromosome. **a:** Conversion times from VCF to other formats. For large datasets, only *vcf.gz* will be the source format (not VCF) because the uncompressed sizes are beyond what fits on typical hard drives. **b**: Time for GRG construction from different file formats with 20 computational CPU threads. **c**: Time to scan each file type (e.g., if all the data were to be loaded into memory). **d**: Time to convert tree sequences to GRGs against varying numbers of samples with the simulated data .

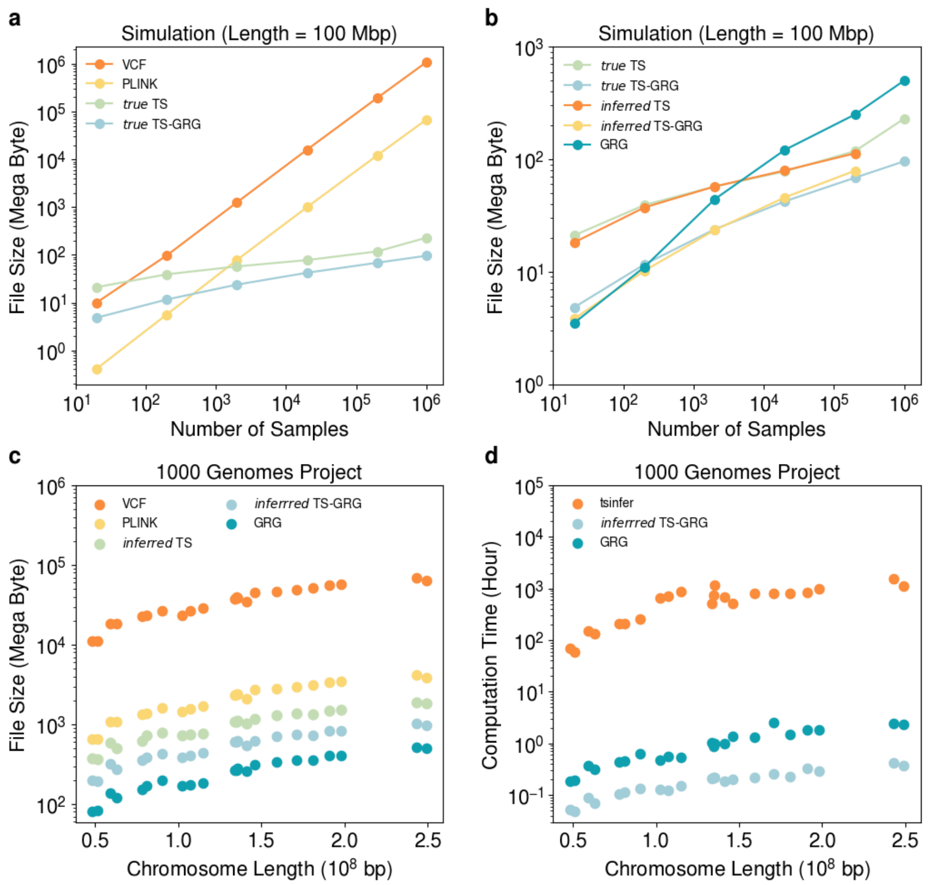


**Figure S7: Investigating the optimality of GRG construction.** A simulated genetic sequence of 100Mbp is used in the benchmark for panels **a** and **b**. **a:** Comparisons of the file sizes among VCFs (VCF), bed files (PLINK), true tree sequences from *msprime* (*true* TS), and GRGs converted from true tree sequences (*true* TS-GRG). **b**: Comparisons of file sizes of *true* TS, *true* TS-GRG, inferred tree sequences from *tsinfer* (*inferred* TS), GRGs converted from inferred tree sequences (*inferred* TS-GRG) and GRG constructed from true genotypes of varied sample sizes (GRG). The phased whole-genome sequences from the Phase III 1000 Genomes Project are used for benchmarking for panels **c** and **d**. **c**: Comparisons of the file sizes using data from the 1000 Genomes Project. Each chromosome is represented as a data point. **d**: Runtime of GRG construction and *tsinfer* inference. The inference time of *tsinfer* is in orange. The construction time of GRG is in teal. The light blue dots show the conversion time from the inferred tree sequences to inferred TS-GRGs.

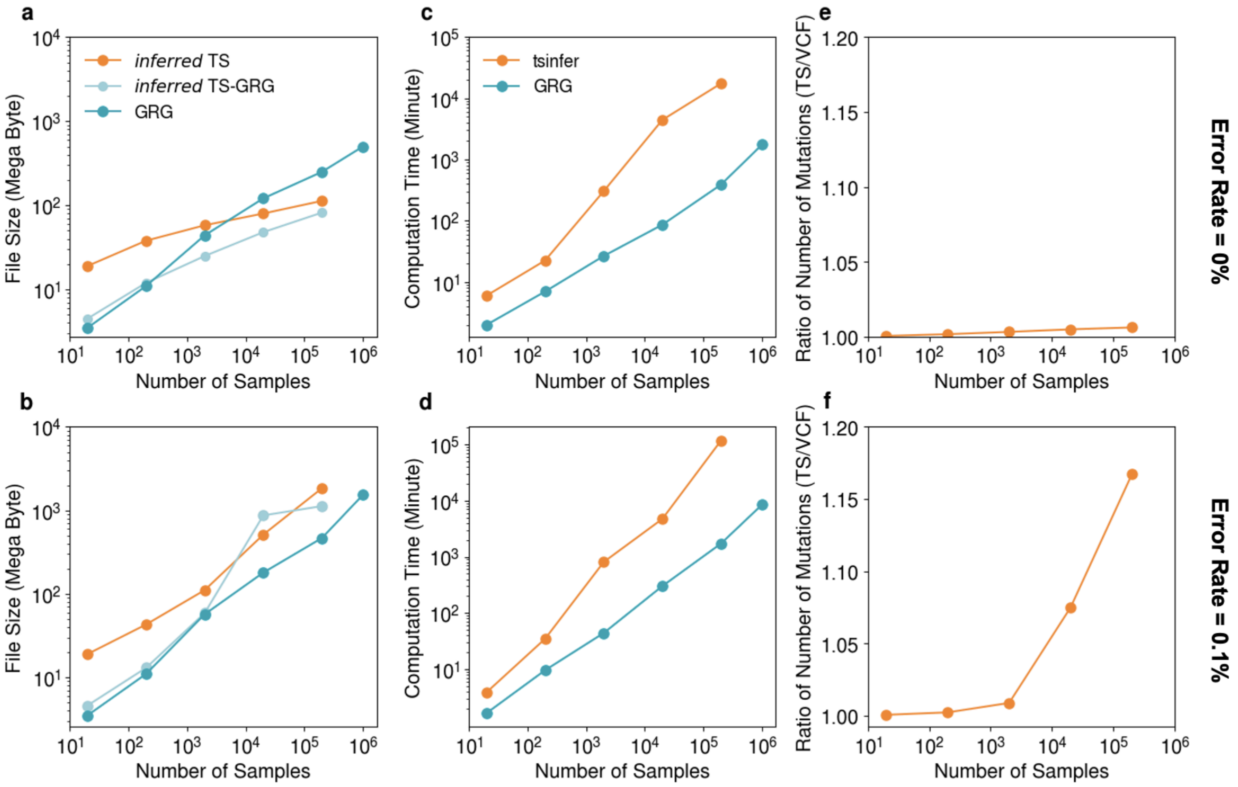


**Figure S8: Robustness of GRG construction with noisy simulated data.** The file sizes (**a-b**) and creation efficiency (**c-d**) of inferred tree sequences and constructed GRGs in simulated data: (**a, c)** without error and (**b, d)** with 0.1% error. The error is introduced by manually flipping individuals' genotypes according to the specified error rate to mimic the sequencing or genotyping error in empirical data. We compare the file sizes and creation times of the inferred tree sequences from *tsinfer* (*inferred* TS), the GRG converted from the inferred TS (*inferred* TS-GRG) and the GRG constructed directly from the genotypes (GRG). **e**, **f:** the ratio of the number of mutations in inferred tree sequences to the number of input alternative alleles in VCF with different error rates in the data.

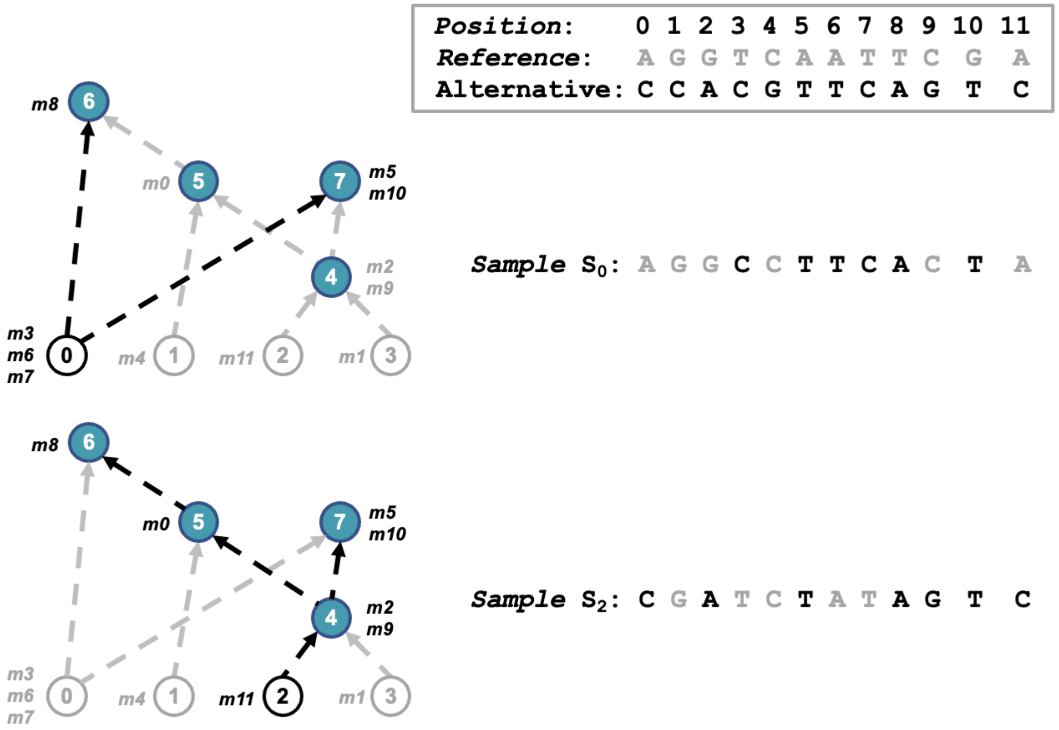


**Figure S9: Querying haplotypes in a GRG.** Here, we present two examples of how to query haplotypes in a GRG (e.g., haplotype0 and haplotype2). To achieve this, we initiate traversal from the sample node that represents the haplotype of interest (e.g. node0 for haplotype0). We then navigate upward from the current node to all parent nodes (e.g., node6 and node7) to collect all mutations occurring in the path (e.g., *m3*, *m6*, *m7*, *m8*, *m5*, *m10*). The other positions without collected mutations are designated as reference alleles.

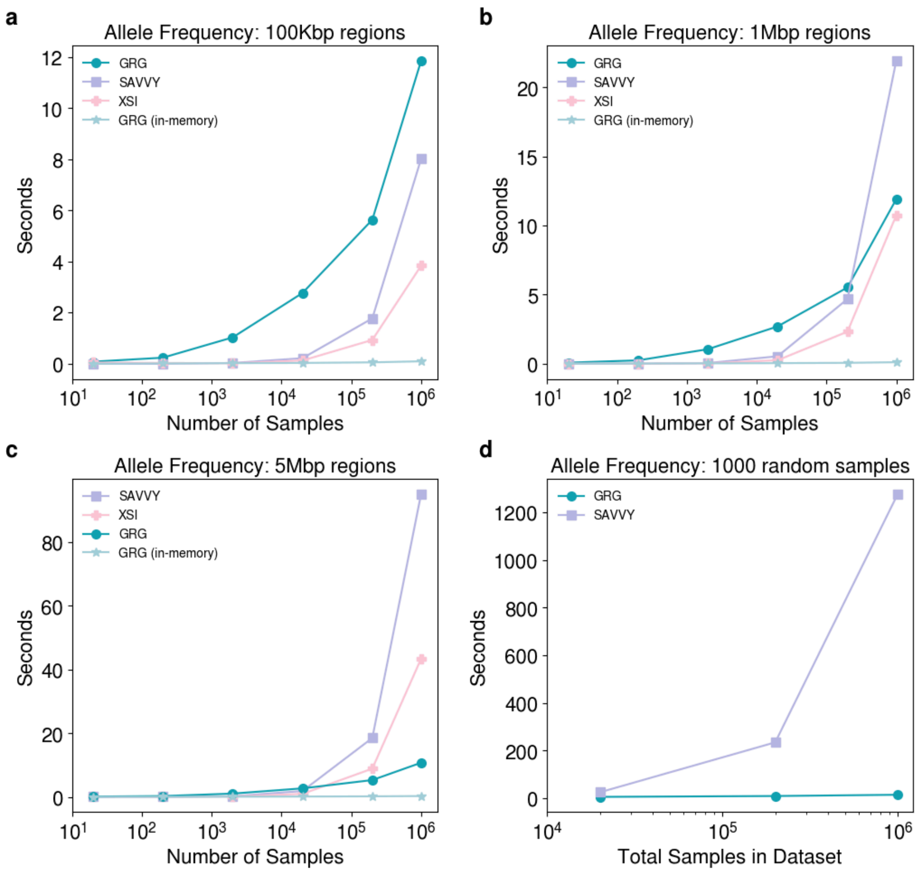


**Figure S10: Computing allele frequency on subsets of the dataset. a**: Average time to compute allele frequency on three separate regions of size 100Kbp. **b:** Average time to compute allele frequency on three separate regions of size 1Mbp. **c:** Average time to compute allele frequency on three separate regions of size 5Mbp. In all three panels (**a**, **b**, **c**) the "GRG (in-memory)" time is the time to compute allele frequency by traversing the graph, once the graph is loaded into RAM. The "GRG" time includes the time to load the graph from disk. **d:** Time to compute allele frequency for a subset of 1000 randomly chosen samples. The same random subset was used for Savvy and GRG.

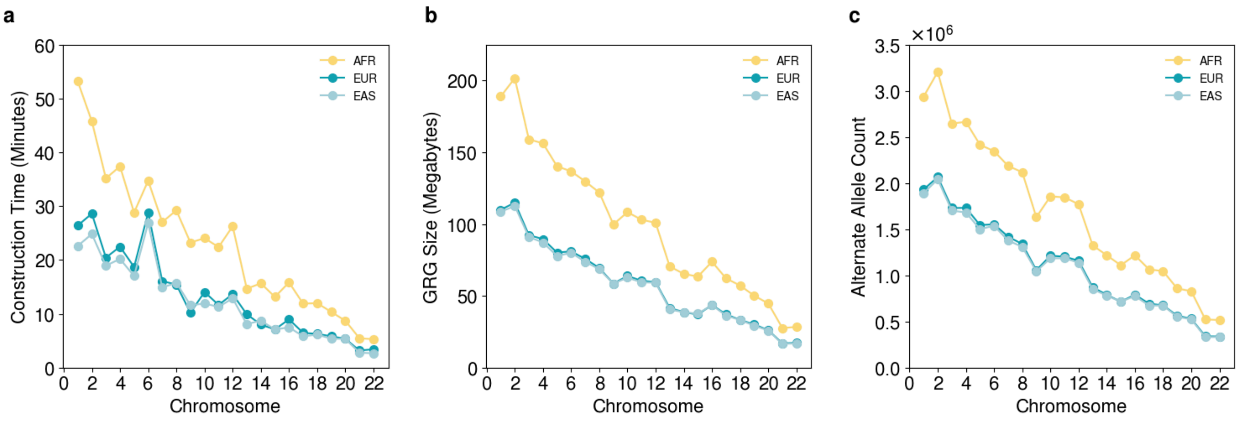


**Figure S11: GRG statistics for three different populations from the 1000 Genomes Project dataset . a**: Time to construct GRG. **b:** GRG file size. **c:** Number of alternate allele values in the dataset. If every site were bi-allelic, this would be equal to the number of sites. GRG explicitly represents every alternate allele value, hence we expect the size (and construction time) to be influenced by the number of alternate alleles.

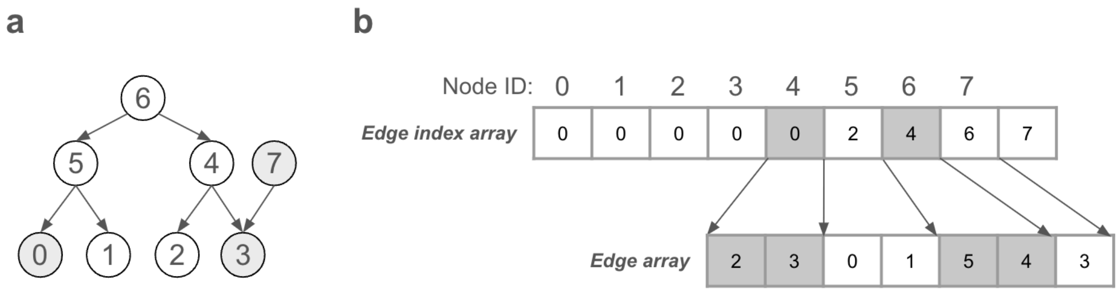


**Figure S12: Compressed Sparse Row (CSR) format of a graph.** **a**: A very simple GRG. **b**: The CSR representation of the GRG contains two arrays: the edge index array and the edge array. The edge index array maps each node ID to its starting position in the edge array. The edge array contains, starting at position *i*, the list of node IDs that are targeted in the direction graph. For example, the first two items in the edge array are 2,3 and indicate that node 4 has edges to nodes 2 and 3.

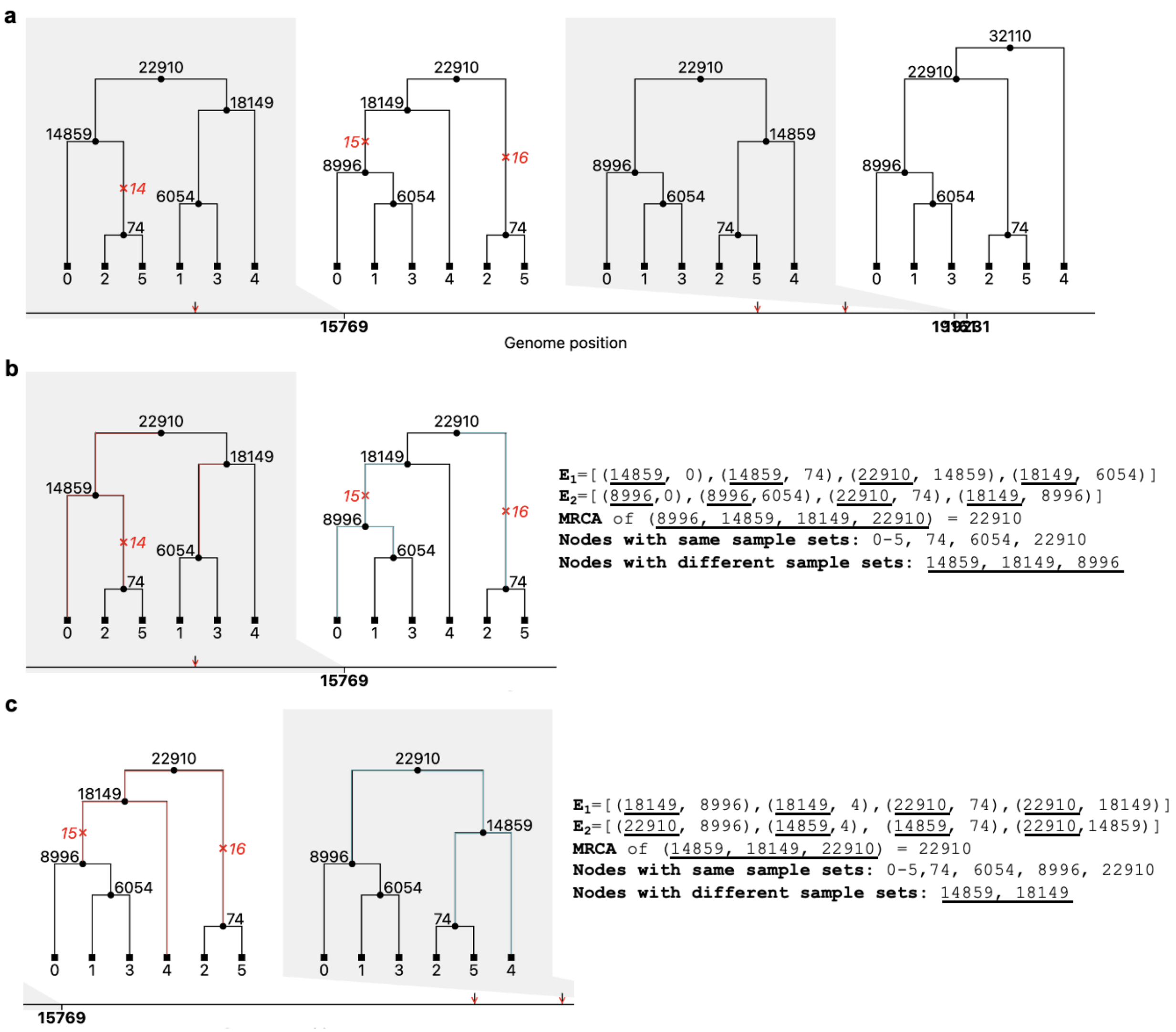


**Figure S13: Converting GRGs from tree sequences. a**: an example of a tree sequence with 4 trees. **b-c**: two recombination events from $\mathcal{T}_i$ to $\mathcal{T}_{i+1}$. $\mathbf{E}_1$ and $\mathbf{E}_2$ represent the list of edges that are deleted (edges in red) or inserted (edges in dark green) from $\mathcal{T}_i$ to $\mathcal{T}_{i+1}$ respectively. The parent nodes of these edges (i.e., nodes that are underscored in the edge lists) potentially have different sample sets in two trees. **b** shows the topological change from $\mathcal{T}_0$ to $\mathcal{T}_1$ and **c** shows the change from $\mathcal{T}_1$ to $\mathcal{T}_2$.

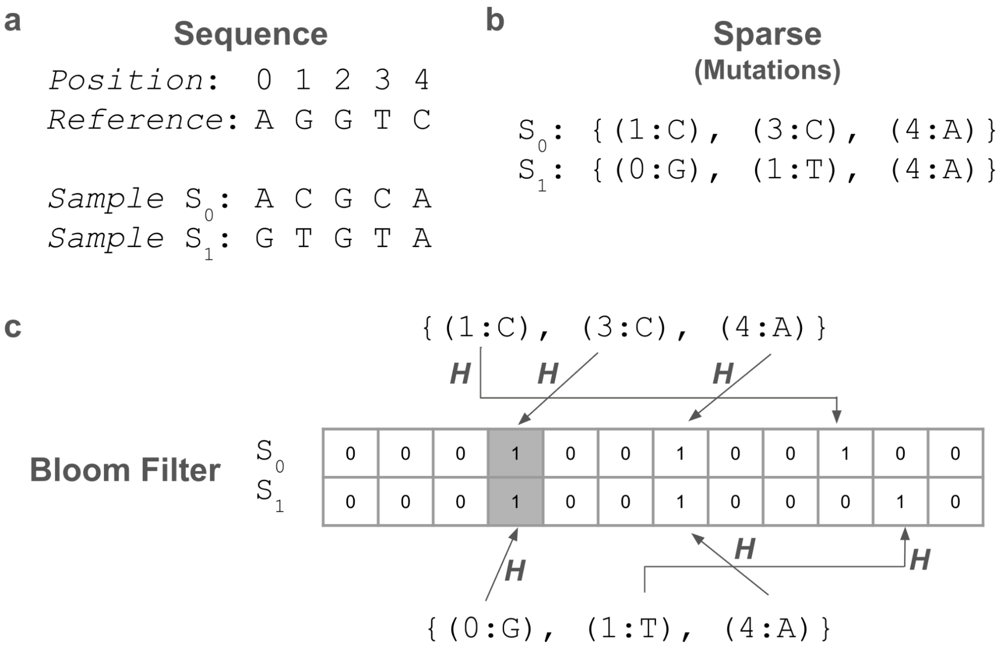


**Figure S14: Three different representations of a haplotype sample. a**: An aligned sequence of nucleotides. **b**: The same sequence as **a**, but represented by lists of (Position:Nucleotide) pairs for *only* the positions where the sample differs from the reference. We call these mutation lists. **c**: The same as **b**, but each mutation is hashed to obtain an index in a bitvector. This is equivalent to a bloom filter with $k = 1$ and $m = 12$. The shaded gray cells demonstrate a false positive: the hash function $H$ maps (3:C) and (0:G) to the same bit, even though they are different mutations.

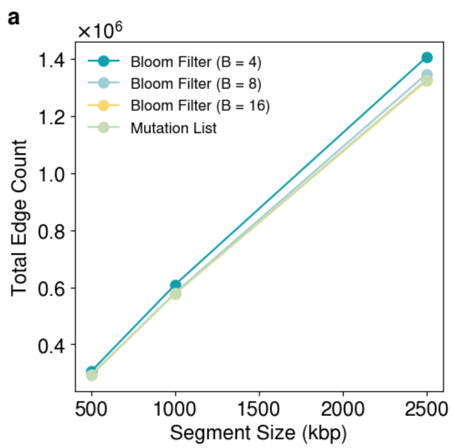


**Figure S15: Edge counts and singleton edge counts for GRGs on a small section of the genome.** The precise haplotype representation (Mutation List) and bloom filter representation with 16-bits per mutation are almost identical in their effect on GRG size. The 4-bit-per-mutation bloom filter is the worst, but only by a very small magnitude.

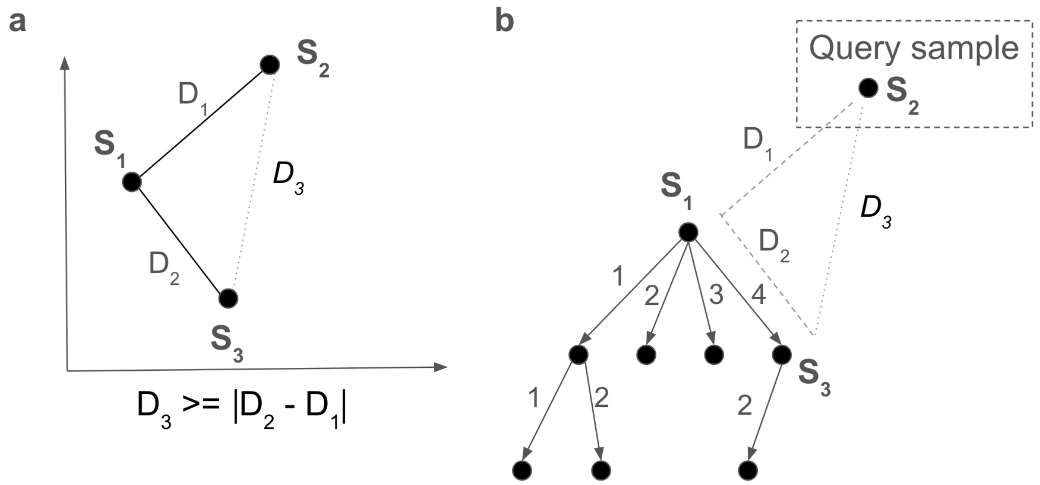


**Figure S16: a**: illustration of the reverse triangle inequality using a triangle in the plane with distances labeled between the three points. Without computing distance $D_3$, we know its lower bound (given $D_2$, $D_1$). The BK-Tree in **b** uses this property to avoid examining entire subtrees. If we are finding the nearest neighbor of $S_2$, and we have already seen a sample with distance $D_{best}$,, we can skip the subtree rooted at $S_3$ if $|D_2 - D_1| >= D_{best}$. We must compute $D_1$ = Hamming($S_2$, $S_1$) during our search, but $D_2 = 4$ is already computed as an edge label.

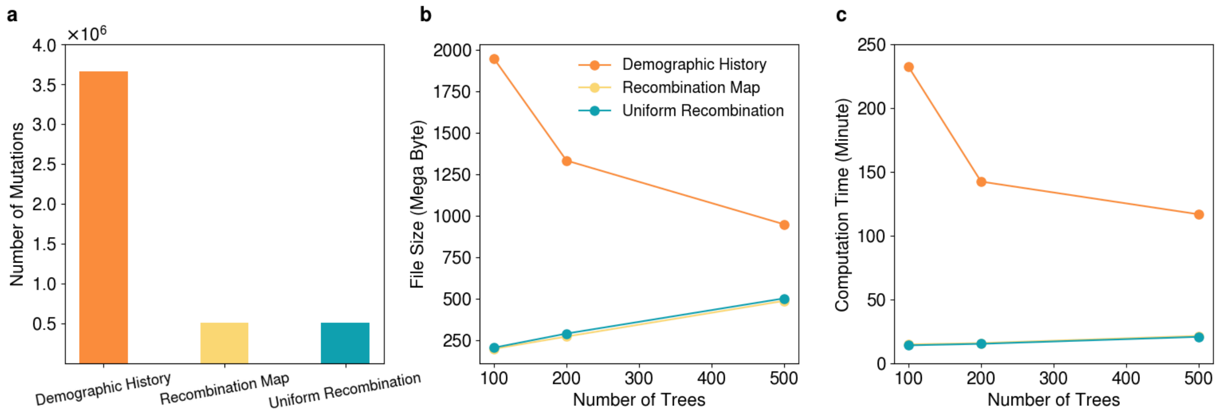


**Figure S17: Constructing GRGs under different simulation settings.** To investigate the impact of simulation scenarios on GRG construction, we tested three different simulation datasets with 100,000 diploid individuals: a flat recombination rate of $10^{-8}$ / (bp $\times$ generation) (teal), a realistic recombination map adapted from the 1000 Genomes Project (utilizing the first 100Mbp of chromosome 1, yellow) and a combination of a realistic recombination map, gene conversions and a demographic model with a four-population out-of-Africa history (orange). **a**: The number of mutations occuring in different simulations with the same simulated generations. **b**: The resulting file size of constructed GRGs with different parameter *T* settings during GRG construction in various simulations. **c**: The construction time for GRGs using different parameter *T* settings.

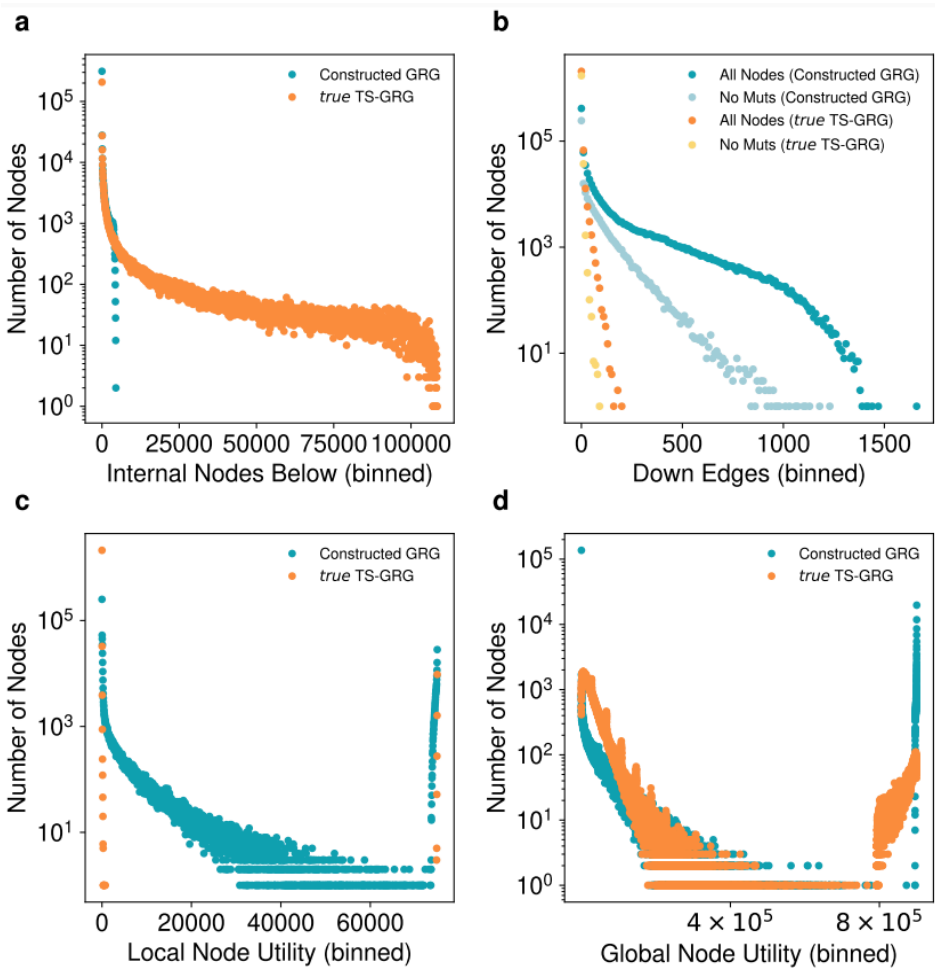


**Figure S18: Histograms of GRG graph statistics for simulated data of length 100Mbp. a**: Distribution (bin size: 10) of the internal nodes beneath each mutation node. More internal nodes beneath a given node is an approximation for the amount of hierarchy below that node. **b**: Distribution (bin size: 50) of nodes with different numbers of down edges. Compares all nodes vs. all non-mutation nodes. **c**: Distribution (bin size: 10) of nodes and their *local utility*. Local utility is just ($Edges_{\mathrm{IN}} \times Edges_{\mathrm{OUT}}$) - ($Edges_{\mathrm{IN}} + Edges_{\mathrm{OUT}}$), i.e. how much savings this node provides in isolation, beyond the sparse matrix representation. **d**: Distribution (bin size: 30) of nodes and their *global utility*. Global utility is the same idea as local utility, but looks at the context of the entire graph: ($Mutations_{\mathrm{ABOVE}} \times Samples_{\mathrm{BENEATH}}$) - ($Edges_{\mathrm{ABOVE}} + Edges_{\mathrm{BENEATH}}$).

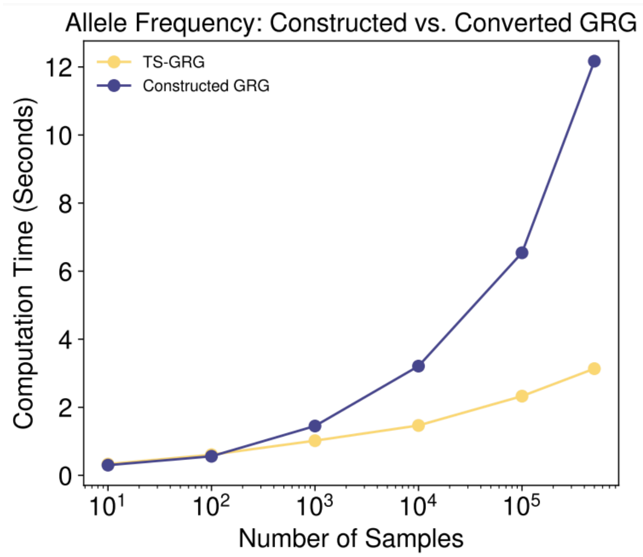


**Figure S19: Computation time for constructed vs. converted GRG.** The time to compute allele frequency using the constructed GRG (from tabular data) compared against the GRG converted from the true tree-sequence. The (true) TS-GRG is more compact than the constructed GRG, hence it is also faster for computation.

Kleinberg, Jon, and Éva Tardos. 2006. *Algorithm Design*. Boston: Pearson/Addison-Wesley.

McVean, Gilean A. T., Simon R. Myers, Sarah Hunt, Panos Deloukas, David R. Bentley, and Peter Donnelly. 2004. “The Fine-Scale Structure of Recombination Rate Variation in the Human Genome.” *Science* 304 (5670): 581–84. https://doi.org/10.1126/science.1092500.

Pratto, Florencia, Kevin Brick, Pavel Khil, Fatima Smagulova, Galina V. Petukhova, and R. Daniel Camerini-Otero. 2014. “Recombination Initiation Maps of Individual Human Genomes.” *Science* 346 (6211): 1256442. https://doi.org/10.1126/science.1256442.

Rivest, R. 1992. “The MD5 Message-Digest Algorithm.” RFC1321. RFC Editor. https://doi.org/10.17487/rfc1321.

The 1000 Genomes Project Consortium, Corresponding authors, Adam Auton, Gonçalo R. Abecasis, Steering committee, David M. Altshuler, Richard M. Durbin, et al. 2015. “A Global Reference for Human Genetic Variation.” *Nature* 526 (7571): 68–74. https://doi.org/10.1038/nature15393.

The International HapMap Consortium. 2007. “A Second Generation Human Haplotype Map of over 3.1 Million SNPs.” *Nature* 449 (7164): 851–61. https://doi.org/10.1038/nature06258.

Wong, Yan, Anastasia Ignatieva, Jere Koskela, Gregor Gorjanc, Anthony W Wohns, and Jerome Kelleher. 2024. “A General and Efficient Representation of Ancestral Recombination Graphs.” Edited by G Coop. *GENETICS*, July, iyae100. https://doi.org/10.1093/genetics/iyae100.

Zhang, Brian C., Arjun Biddanda, Árni Freyr Gunnarsson, Fergus Cooper, and Pier Francesco Palamara. 2023. “Biobank-Scale Inference of Ancestral Recombination Graphs Enables Genealogical Analysis of Complex Traits.” *Nature Genetics* 55 (5): 768–76. https://doi.org/10.1038/s41588-023-01379-x.
