## Supplementary Table for "Genotype Representation Graphs: Enabling Efficient Analysis of Biobank-Scale Data"

### SUPPLEMENTAL TABLE

**Data Availability**

GISAID Identifier: EPI_SET_230329cd
doi: 10.55876/gis8.230329cd

All genome sequences and associated metadata in this dataset are published in GISAID's EpiCoV database. To view the contributors of each individual sequence with details such as accession number, Virus name, Collection date, Originating Lab and Submitting Lab and the list of Authors, visit 10.55876/gis8.230329cd

**Data Snapshot**

- EPI_SET_230329cd is composed of 11,126,604 individual genome sequences.
- The collection dates range from 2020-01-01 to 2022-07-04;
- Data were collected in 218 countries and territories;
- All sequences in this dataset are compared relative to hCoV-19/Wuhan/WIV04/2019 (WIV04), the official reference sequence employed by GISAID (EPI_ISL_402124). Learn more at https://gisaid.org/WIV04.
